# The Marginal Cells of the *Caenorhabditis elegans* Pharynx Scavenge Cholesterol and Other Hydrophobic Small Molecules

**DOI:** 10.1101/565473

**Authors:** Muntasir Kamal, Houtan Moshiri, Lilia Magomedova, Duhyun Han, Ken CQ Nguyen, May Yeo, Jess Knox, Rachel Bagg, Amy M. Won, Karolina Szlapa, Christopher Yip, Carolyn L. Cummins, David H. Hall, Peter J. Roy

## Abstract

The nematode worm *Caenorhabditis elegans* is a bacterivore filter feeder. Through the contraction of the worm’s pharynx, a bacterial suspension is sucked into the pharynx’s lumen. Excess liquid is then shunted out of the buccal cavity through ancillary channels that are made from specialized pharyngeal cells called marginal cells. Through the characterization of our library of worm-bioactive small molecules (*a.k.a.* wactives), we found that more than one third of wactives visibly accumulate inside of the marginal cells as crystals or globular spheres. Wactives that visibly accumulate are typically more hydrophobic than those that do not. To understand why wactives accumulate specifically in marginal cells, we performed a forward genetic screen for mutants that resist the lethality associated with one crystallizing wactive. We identified a presumptive sphingomyelin-synthesis pathway that is necessary for crystal and sphere accumulation. Sphingomyelin is a phospholipid that is enriched in the outer leaflet of the plasma membranes of most metazoans. We find that the predicted terminal enzyme of this pathway, sphingomyelin synthase 5 (SMS-5), is expressed in the pharynx, contributes to sphingomyelin abundance, and that its expression in marginal cells is sufficient for wactive accumulation. We also find that the expression of SMS-5 in the marginal cells is necessary for the proper absorption of exogenous cholesterol, without which *C. elegans* cannot develop. We conclude that the sphingomyelin-rich plasma membrane of the marginal cells acts as a sink to scavenge important hydrophobic nutrients from the filtered liquid that might otherwise be shunted back into the environment.

**One sentence summary:** The anterior pharynx of *C. elegans* is a Sink for Hydrophobic Small Molecules

## Introduction

The survival of life in the wild is dependent on enduring the boom and bust cycles of nutrient availability. This is especially true for the free-living nematode *Caenorhabditis elegans* that must survive extreme population bottlenecks because of limited nutrients [1]. Under ideal conditions, *C. elegans* will undergo embryogenesis and develop through four larval stages called L1 through L4 to reach reproductive maturity in as little as three days [2]. Upon nutrient deprivation, over-crowding, or excessive temperature, the animal will enter an alternative stress-resistant long-lived dispersal state called dauer instead of L3 [3]. Extensive sampling of nematodes in the wild reveals that *C. elegans* is most often found in the dauer state, unlike other free-living nematodes outside of the *Caenorhabditis* genus that are found to be actively proliferating [1]. Hence, *Caenorhabditis* seems particularly sensitive to stress, including nutrient deprivation.

*C. elegans* is a sterol auxotroph [4]. In the laboratory, *C. elegans’* food (*E. coli*) is supplemented with cholesterol, but other select fungal, plant and animal sterols will also suffice [5]. The worm modifies the sterols for use in signaling, and unlike vertebrates, may be a structural component within the membranes of only a limited number of cells [4]. Depending on the extent and timing of sterol withdrawal, *C. elegans* will either enter the dauer state or fail to develop and perish [5]. A lack of select sterols, which are relatively insoluble and rare in the wild, is likely one factor that contributes to *Caenorhabditis* existing in the dauer state outside of the laboratory [6]. Mechanisms that maximize the absorption of these precious nutrients must therefore be essential to the overall success of the species.

*Caenorhabditis* is a filter-feeder [7, 8]. Its feeding organ, called the pharynx, has a three-fold symmetry with three muscles divided by three sets of ‘marginal cells’, all radially-oriented with respect to the central lumen (see figures for schematics) [7]. The first step of the feeding cycle is that the radially aligned muscles of the pharynx contract, thereby opening the central lumen and creating negative pressure that sucks the bacterial suspension from the environment into the lumen. The pharynx then relaxes in an anterior to posterior wave, concentrating the suspended bacterial cells centrally within the lumen. Because of increasing pressure, the excess liquid escapes to the environment in a posterior-to-anterior manner via channels within the marginal cells [9]. The microbes are then macerated in the posterior pharynx and passed on to the neighboring intestine via peristaltic movements [7].

Through a survey of how *C. elegans* interacts with worm-active (wactive) synthetic small molecules, we discovered that relatively hydrophobic molecules visibly accumulate in the marginal cells of the anterior pharynx. No other tissue visibly accumulates hydrophobic small molecules. We identified several components of a presumptive sphingomyelin synthesis pathway, at least one of which is specifically expressed in the pharynx, that are necessary for this accumulation. We find that the expression of the presumptive terminal enzyme of this pathway (SMS-5) specifically in the marginal cells is sufficient for wactive accumulation. These observations led us to the finding that the marginal cells play an important role in the absorption of cholesterol, which is consistent with previous work showing that the pharynx accumulates sterols and other natural products [10, 11]. Together, our observations indicate that the marginal cells scavenge precious nutrients from fluid that would otherwise be discarded by the worm. This nutrient-scavenging mechanism may be important in allowing *Caenorhabditis* to survive periods of limited nutrient availability.

## Results

### Select Wactive Compounds Accumulate as Crystals and Globular Spheres in the Anterior Pharynx

Observations from our previous drug screens revealed that animals incubated in some compounds have unusually dark pharynxes when visualized at 100X magnification [12-14]. We investigated this phenomenon further by incubating synchronized first larval stage (L1) wild type worms in 238 compounds from our wactive library at a concentration of 30 μM for 48 hours (Supplemental Dataset 1). We visualized the worms at 400X magnification using differential interference contrast (DIC), which allows clear visualization of unusual objects, and polarized light (birefringent) microscopy, which allows for the detection of objects containing molecules that are arrayed in a regular pattern (i.e. are crystalline) [15]. We found that 38 (16%) wactives accumulated as birefringent crystals, and another 33 (14%) accumulated as non-birefringent globular spheres (henceforth referred to as spheres) (Fig 1a-1c). The only cells in which we found these unusual objects are within the corpus (anterior half) of the pharynx.

**Figure 1.**
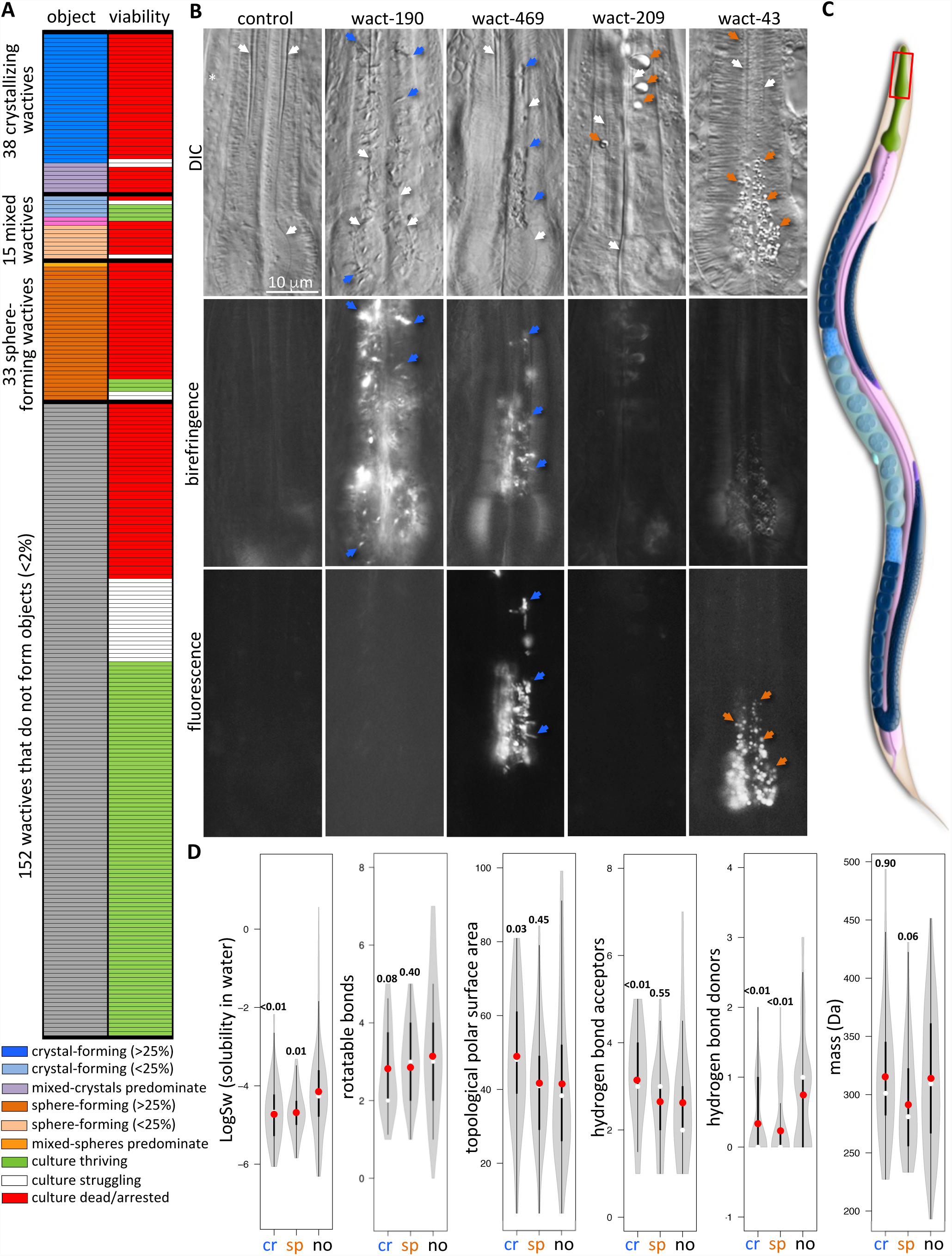
Small Molecules can Crystallize and/or form Globular Spheres in the Marginal Cells of the Anterior Pharynx. **A.** A chart showing our survey of the effects of 238 wactives on *C. elegans* L1 animals. Each row is a distinct wactive. The first column indicates whether a wactive accumulates as a crystal, sphere, or neither at a concentration of 30 μM. The second column indicates whether the wactive also kills/arrests worm development at a concentration of 30 μM. The percentages in the legend refer to the percentage of animals harbouring the indicated objects. See methods for details. **B.** Examples of the accumulation of crystals and spheres in the anterior pharynx of young adults treated for 24 hours to allow easy visualization of the channels. The first row shows differential interference contrast (DIC) images; the second row shows whether the objects are birefringent (and therefore crystalline); and the third row shows whether the object fluoresces. White arrows indicate the channels, blue arrows indicate crystals, orange arrows indicate spheres. The white asterisk indicates where the micrograph was spliced to ensure channel is in the correct focal plane. The scale is the same for all panels. **C.** A schematic of *C. elegans* (courtesy of WormAtlas [40]). The anterior pharynx is boxed. **D.** Violin plots of the physical-chemical properties of molecules that crystalize (cr), form spheres (sp), or fail to form objects in the animal (no). The gray shape represents all results and the thickness indicates how common that value is within the dataset; a white circle indicates the median; a red circle indicates the mean; the centre thick line represents one standard of deviation; the thinner line represents the second standard of deviation. The number above each plot shows the significance of the difference compared to the ‘no-object’ data using the nonparametric two-tailed Mann-Whitney U test.

We investigated whether the crystals and spheres are a response of the worm to the wactives or whether the objects are likely to be the wactives themselves. We found that one of the crystallizing wactives, wact-469 fluoresces yellow-green (Supplemental Fig 1), while no other crystallizing wactive that we investigated in detail fluoresces using our detection methods. We found that crystals that form in response to wact-469 fluoresce yellow-green, but crystals that form in response to other wactives like wact-190 do not fluoresce (Fig 1b). Similarly, the wactive wact-43 fluoresces blue, and results in blue fluorescent spheres in the animal, whereas other non-fluorescent wactives like wact-209 do not produce fluorescent spheres (Fig 1b). We conclude that the crystals and spheres are at least partly, if not entirely, composed of the respective wactive compound.

We examined some basic physicochemical features of the wactives to better understand the properties associated with crystallization and sphere formation. We found that both crystal-forming and sphere-forming wactives are less hydrophilic (and have fewer hydrogen bond donors, which is related to hydrophilicity) than the wactives that do not generate unusual objects (*p*≤0.01) (Fig 1d). This finding is consistent with the molecules precipitating out of solution when concentrated within the animal. One feature that distinguishes the crystal-forming compounds from those that form spheres is that the former have a greater topological polar surface area (*p*<0.05) (Fig 1d). A key determinant of whether a membrane protein will crystallize in x-ray crystallography studies is its topological polar surface area; more exposed polar groups increase the likelihood that a membrane protein will crystalize easily [16]. This raises the possibility that the crystallizing wactives are associating with the plasma membrane and coming out of solution in crystalline form due to their relatively large polar surface area.

### Wactive Crystals May Contribute to the Developmental Arrest and/or Death of ***C. elegans***

The majority of wactives that form crystals and spheres at 30 μM also induce developmental arrest and/or death of L1 larvae at that concentration (Fig 1a). Of note, for many object-forming wactives, we find day-to-day variability in whether they outright kill the L1 culture (resulting in disintegrated animals) or simply lead to persistent developmentally-arrested L1s.

To better understand the relationship between object formation and lethality/arrest, we performed a dose-response analysis on a small subset of wactives that form crystals or spheres (Fig 2a). We collected synchronized first larval stage animals (L1s) and inspected the pharynxes and other tissues at higher magnification after 48 hours of incubation in the wactives. In parallel, we performed our standard six day viability assay [14]. We found a strong correlation between the presence of objects in the anterior pharynx and the associated lethality/arrest (an average R^2^ of 0.964).

**Figure 2.**
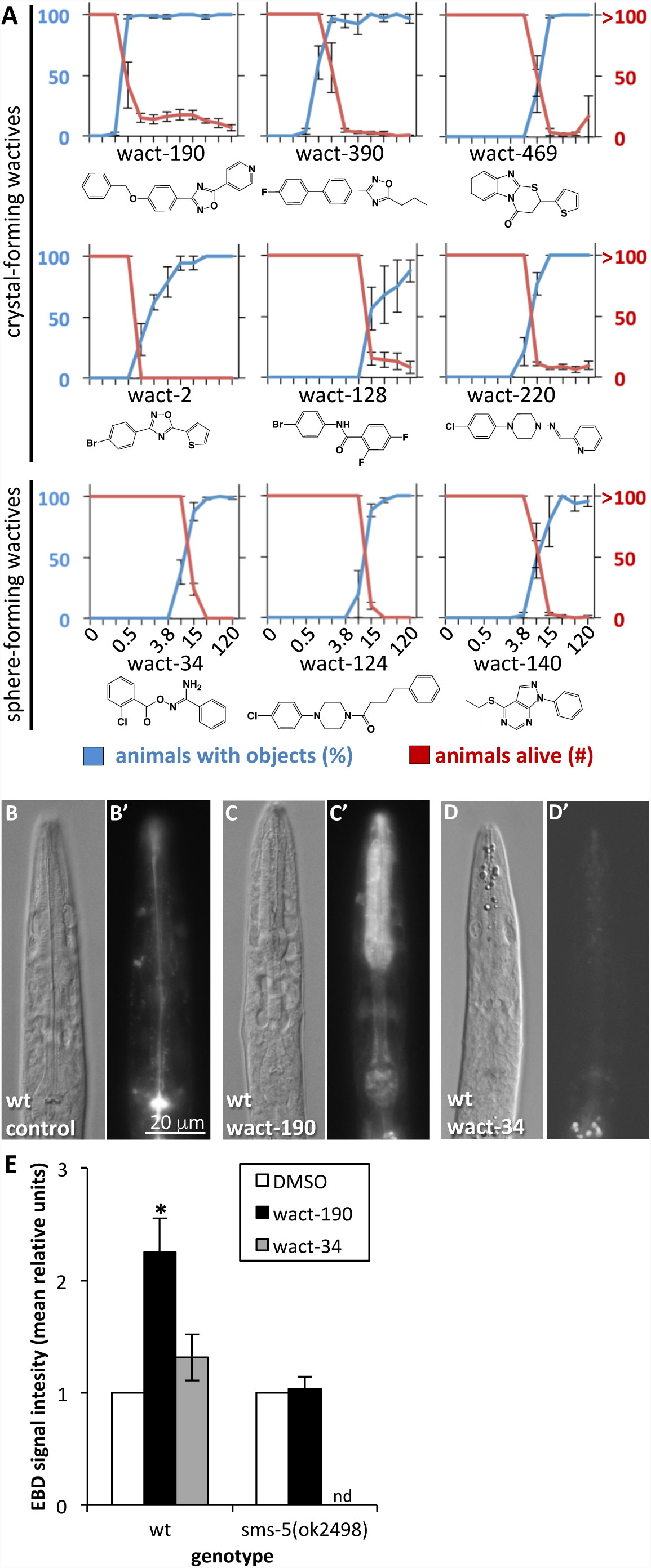
The Dose-Response Relationship Between the Lethality and Object Formation Induced by Select Wactives. **A.** Crystal and sphere formation in the pharynx was analyzed after a 48 hour incubation of L1 larvae in the respective wactive and is indicated in blue. The lethality induced by the respective wactive, assayed after 50 L1s were incubated with the molecules for six days, is indicated in red. Animals in each well were counted if not over-grown with worms; otherwise, the well was scored as 100 animals. Low numbers of animals (see wact-190 or wact-128 for example) are invariably arrested L1s. See methods for details. The two-fold serial dilution dose of the wactive, in micromolar, is indicated at the bottom of the three columns of graphs. The small molecule structure is illustrated below each graph. **B-E.** An analysis of the ability of Evans Blue dye (EBD) to penetrate the anterior pharynx. L1-stage worms were incubated with control (1% DMSO only) or 60 µM of the indicated compound for 24 hours. Worms were then incubated in EBD for 4 hours. **B-D.** We see an enrichment of EBD signal in the anterior pharynx of worms treated with the crystal-forming wact-190, but not the sphere-forming wact-34 or control. The scale is the same for all panels. **E.** Quantification of the EBD signal intensity of wact-190-treated worms relative to the DMSO negative control treated worms. See methods for details. Asterisk indicates a significant difference between the control and the wact-190 treated sample (*p*<0.05). Error bars represent standard error of the mean (SEM).

Many of the object-forming wactives are structurally diverse but have crystal- or sphere-formation in common (Fig 2a). These observations raised the possibility that the objects themselves are contributing to the death or arrest of the animal. We reasoned that the crystals might perforate the membrane of the anterior pharynx and lead to death. We have no similar hypothesis about the spheres. We incubated the animals in the crystal-forming wactive wact-190, and sphere forming wact-34, and tested whether a membrane impermeable dye permeates the pharynx. We found that wact-190-exposed worms allow the permeation of Evans Blue dye [17] (Fig 2b, 2c, and 2e). By contrast, wact-34 failed to allow Evans Blue dye entry (Fig 2d and 2e). These data are consistent with the idea that crystal formation may contribute to the death or arrest of the nematode by disrupting plasma membrane integrity.

### The Marginal Cells of the Anterior Pharynx Act as a Sink for Select Wactives

To investigate the dynamics of crystal formation, we performed a time-course analysis of wact-190 crystal formation. We found that small birefringent objects can be seen as early as 30 minutes after incubation with wact-190 (Fig 3a-3c). By 48 hours, the pharynx lumen of many animals is entirely filled with wact-190 crystals.

**Figure 3.**
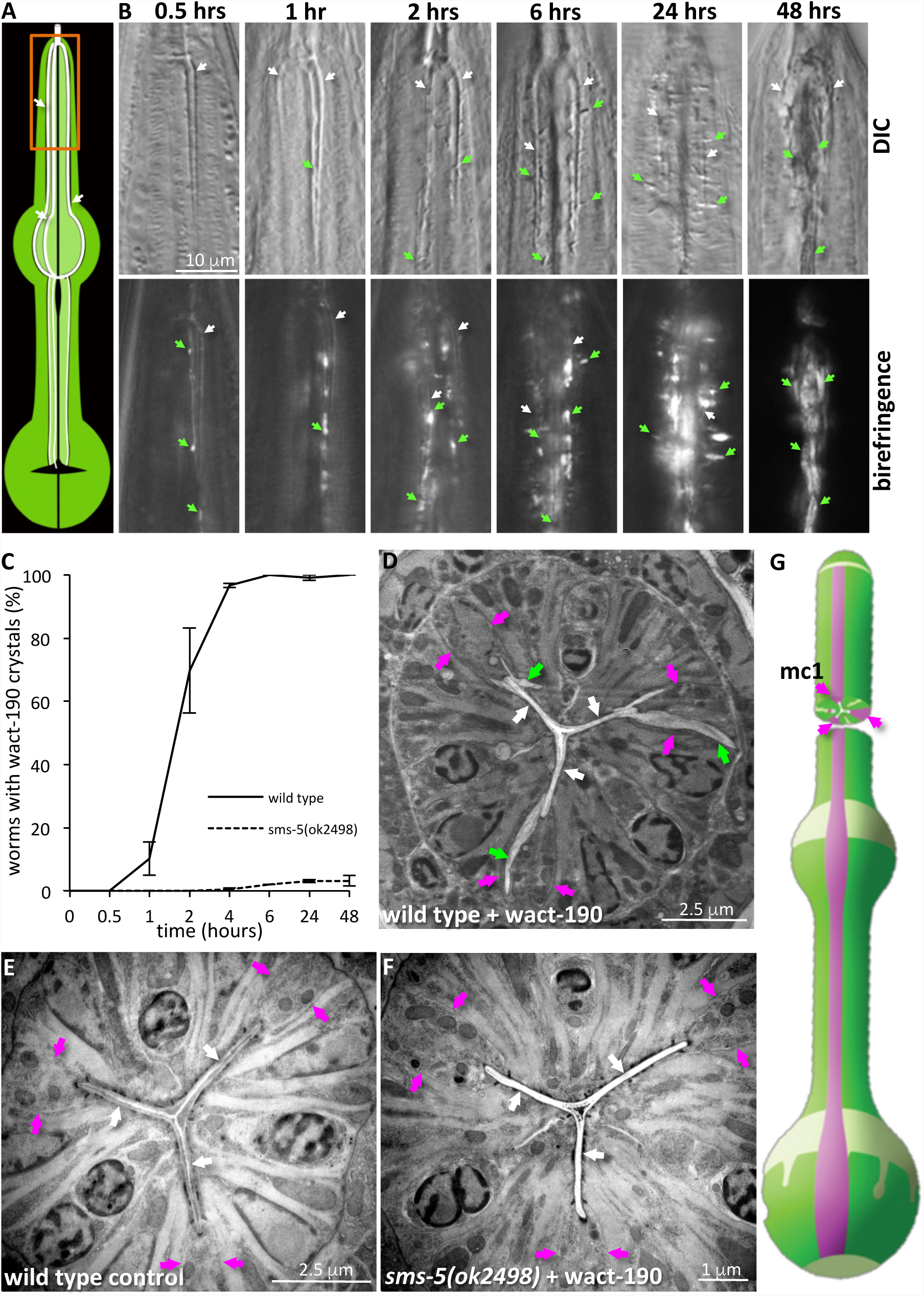
A Microscopic Analysis of wact-190 Crystal Formation. **A.** A schematic of the pharynx with the three channels indicated with white arrows; the central lumen and grinder are in black. The procorpus of the anterior pharynx, which is shown in ‘**B**’, is boxed in orange. **B.** Qualitative analysis of wact-190 crystal formation over time. Synchronized forth stage (L4) animals were grown in 60 μM of wact-190 in liquid culture for the time indicated at the top of each column. The top images show DIC micrographs; the corresponding bottom images are taken with polarized filters to visualize birefringence. Each pair of images shows the procorpus (anterior quarter) of the pharynx. The white arrows indicate channels; green arrows indicate select crystals. The scale is the same for all panels. **C.** Quantitative analysis of wact-190 crystal formation over time. Synchronized first stage (L1) worms of the indicated genotype were incubated in liquid with 60 μM wact-190 for the indicated time period and crystals were identified by their birefringence. **D-F**. Transmission electron micrographs (TEMs) of pharynx cross-sections of the indicated genotype and wactive-treatment. The channels are indicated with white arrows; presumptive crystals with green arrows; marginal cell plasma membranes with fuchsia arrows. See Supplemental Fig 3 for transverse sections. **G.** A schematic of the pharynx illustrating the approximate area of the cross sections imaged by TEM. The marginal cells are indicated in fuchsia. Image adapted from WormAtlas with permission.

Early in the time-course, wact-190 crystals accumulate near the channels of the anterior pharynx (Fig 3a-3c; also see Fig 1). The channels are formed by specialized cells called marginal cells and are part of the filtering mechanism by which the animal feeds on bacterial cells within its environment (see the introduction). The area of crystal and sphere formation, together with the animals’ filter feeding strategy, raised the possibility that crystals might be filtered from the media, get trapped in the channels, and seed the growth of larger crystals. Two observations argue against this. First, the birefringent crystalline objects that can been seen early in the time course experiment are not found in the lumen of the channels, but are associated with its plasma membrane or extra-cellular matrix (see the 30 minute time point in Figure 3). Second, we found that crystals accumulate in the pharynx of young larvae that hatch within the parent’s uterus and are not exposed to the external media (Supplemental Fig 2). Together, these observations argue against the idea that crystals are seeded from crystals in the media. Instead, it is more likely that soluble small molecule accumulates in the tissue of the anterior pharynx, precipitates out of solution and forms crystals or spheres therein.

To examine the location of crystals in greater detail, we inspected the location of crystals using transmission electron microscopy. In cross-sections of wild type animals incubated with wact-190, we found unusual structures that are consistent with the crystals we see with light microscopy. The unusual objects are located in association with the plasma membrane of the marginal cells and extend into the cytoplasm (Fig 3d-3g). In transverse sections of wild type animals incubated with wact-190, we also observe crystal-like objects emanating from the marginal cells. In addition, we see areas of clear material and/or separation of the plasma membrane from the luminal cuticle of the marginal cells (Supplemental Fig 3). These observations are consistent with the idea that the marginal cells have unique properties that facilitate the absorption of small molecules with the physicochemical properties described above.

### SMS-5 Functions in the Anterior Pharynx to Confer Sensitivity to wact-190 Crystal Formation and Lethality

To better understand the accumulation of crystals in the anterior pharynx, we performed a forward genetic screen to isolate mutants that resist the lethal effects of the crystal-forming wact-190. We reasoned that if crystal formation is related to wact-190’s lethality, mutants that resist its lethality would also resist crystal formation. We screened 1.3 million randomly mutagenized wild type F2 genomes and isolated 46 mutants that resist the lethal effects of wact-190 (Table 1; Supplemental Dataset 2). A screen of 200,000 mutant F1 genomes did not yield wact-190-resistant mutants, suggesting that the resistant mutants isolated in the F2 screens are likely recessive.

**Table 1.**
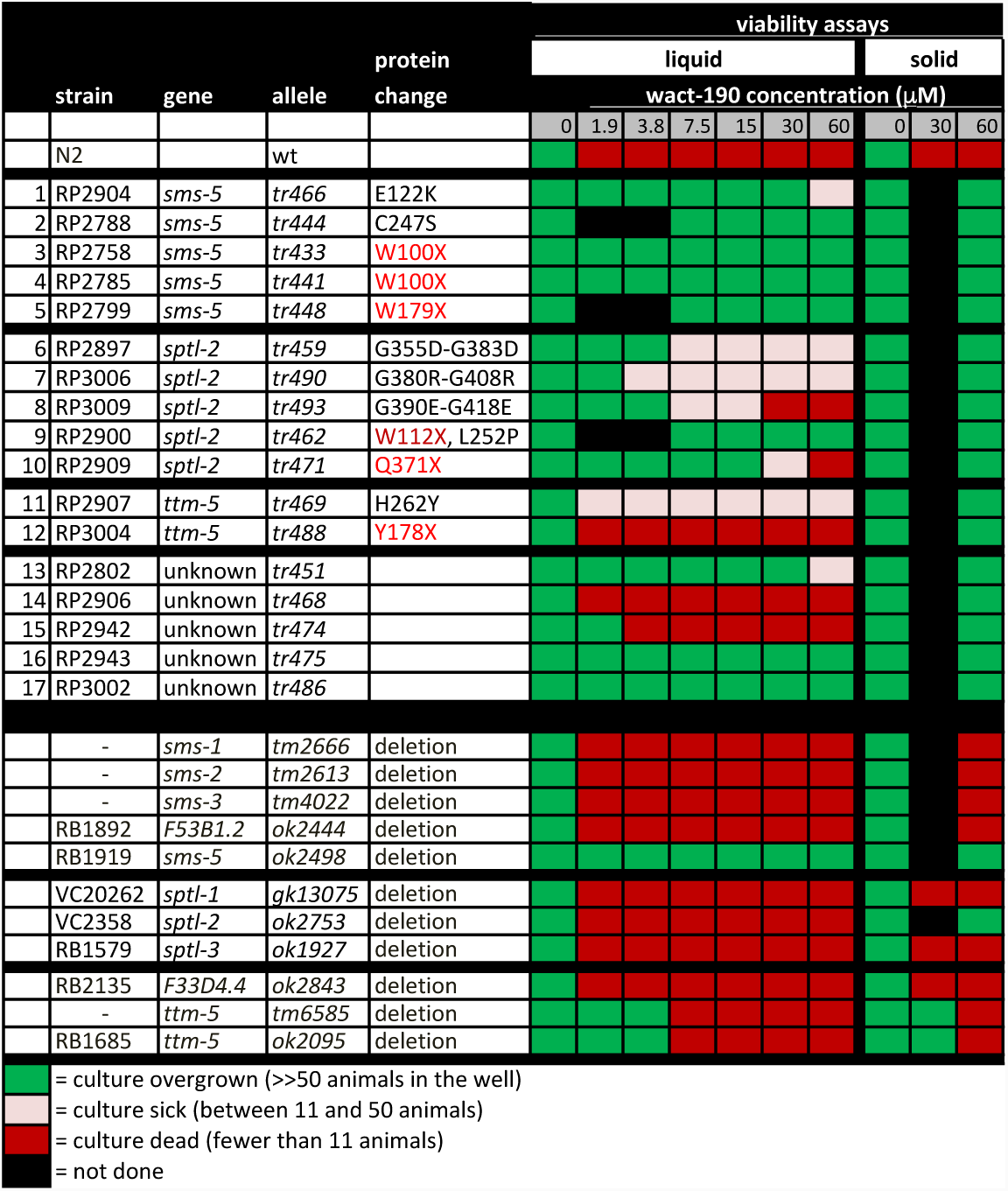
The genes identified in a screen for mutants that resist the lethality induced by of wact-190. Mutants listed in rows 1-17 were isolated in our forward genetic screen for those that resist the lethal/arrest phenotype of wact-190. In addition to these 17, we identified 29 other strains with a mutant allele of *pgp-14* that will be described elsewhere. Additional mutants beyond row 17 are deletion alleles obtained from either the *C. elegans Genetics Centre* or from Shohei Mitani. While all of the mutants isolated in our screen resist wact-190 on solid substrate, only some also resist wact-190 effects in liquid culture. *F53B1.2* is a paralog of the *sms* genes, and *F33D4.4* is a paralog of *ttm-5*. Viability assays were done with at least four replicates (see methods for details). Residue numbers for both isoforms of *sptl-2* are shown. Protein changes noted in red indicate a presumptive null allele; ‘X’ indicates an early non-sense codon. For five wact-190 resistant strains (rows 13-17), the mutant gene responsible for wact-190 resistance has not been identified and are not discussed further.

Sequencing the genomes of the 46 mutants revealed 29 alleles of *pgp-14,* whose contribution to small molecule accumulation will be discussed elsewhere. Sequencing also revealed twelve strains that have mutations in three different components of a predicted *de novo* sphingomyelin biosynthetic pathway (Fig 4a). Sphingomyelin is one of four major phospholipids that constitute the metazoan plasma membrane. Sphingomyelin and phosphatidylcholine are the major lipids in the outer leaflet of the plasma membrane, while phosphatidylserine and phosphatidylethanolamine are the major lipids within the inner leaflet [18, 19].

**Figure 4.**
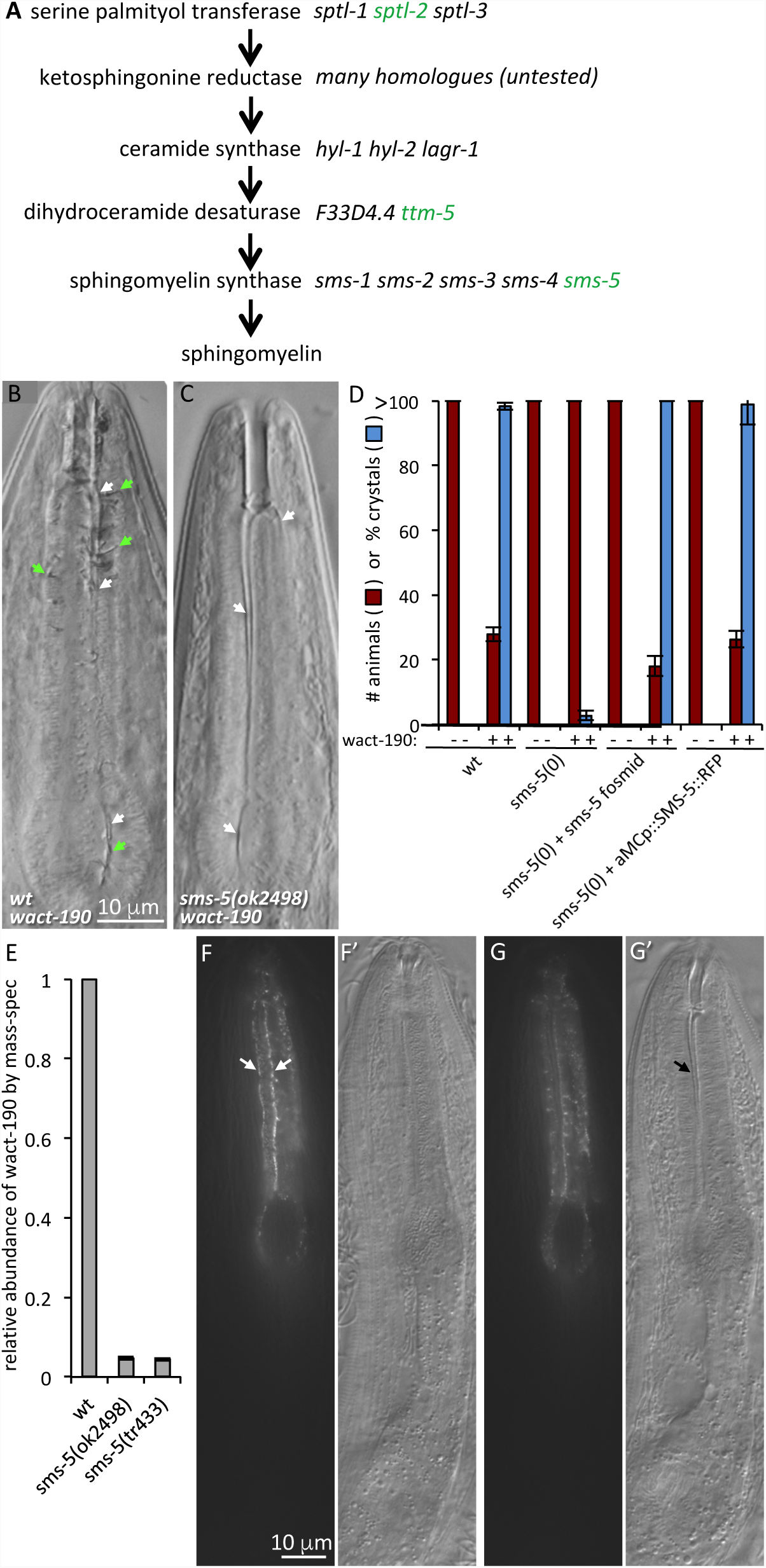
Mutations in a Predicted Sphingomyelin Synthesis Pathway Confer Resistance to wact-190. **A.** A schematic of a predicted sphingomyelin synthesis pathway. Whole-genome sequence analysis indicated that mutations in three genes predicted to play a role in sphingomyelin synthesis (*sptl-2, ttm-5*, and *sms-5*) are sufficient to confer resistance to the lethality induced by wact-190 (see Table 1). Independently-derived deletion alleles were tested for each of these genes (in green), together with their respective paralogs (in black) and only the deletions of *sptl-2, ttm-5*, and *sms-5* confer resistance (see Table 1). **B-C.** 60 μM wact-190 forms crystals in wild type worms but not in the *sms-5(ok2498)* deletion mutant. Animals were incubated in the small molecule for 24 hours as L4 staged animals. Anterior is up; white and green arrows indicate channels and crystals, respectively. **D.** Quantification of wact-190’s effects on viability and crystal formation on the indicated genotype. The *sms-5*-containing fosmid (WRM0626dC03) is harboured as an extra-chromosomal array. The construct expressing SMS-5::RFP from an anterior marginal cell-specific promoter (aMCp::SMS-5::RFP) (see Supplemental Figure 5) is harboured as an integrated transgenic array (*trIs104*). Viability is quantified in a six-day viability assay as described in the methods section. In samples where viability is indicated as <40, the respective wells have only the original larvae in the wells that remain arrested or dead as young larvae. Standard error of the mean is shown. **E.** Accumulation of the indicated small molecule in *sms-5* mutants relative to wild type animals as measured by mass-spectrometry (see methods). Standard error of the mean is shown. **F-G.** Two different focal planes of an adult animal mosaic for the extra-chromosomal (Ex) array harbouring pPRHM1051, a construct consisting of a genomic copy of SMS-5 (fosmid WRM0626dC03) with YFP coding sequence inserted in frame at the C-terminus of SMS-5. The YFP-fusion protein is localized to the anterior marginal cells in this animal and is enriched basally at the borders with adjacent muscle cells (white arrows). See Supplemental Figure 4 for the non-mosaic expression pattern of SMS-5. The scale is the same for all corresponding panels.

Five of the wact-190-resistant strains have a mutation (including three early non-sense alleles) in *sms-5* (*a.k.a.* W07E6.3), which is a gene predicted to encode an integral membrane sphingomyelin synthase and the terminal enzyme in the pathway [20] (Table 1; Fig 4a). Sphingomyelin synthase catalyzes the production of sphingomyelin from phosphatidylcholine and ceramide on the outer leaflet of the plasma membrane [21]. Another five wact-190-resistant strains have a mutation (including two nonsense alleles) in *sptl-2*, which encodes a serine palmitoyltransferase. Two strains have a mutation (including one nonsense allele) in *ttm-5*, which is predicted to encode a sphingolipid delta(4)-desaturase. Both SPTL-2 and TTM-5 are predicted to function upstream of SMS-5 in the production of sphingomyelin [21].

At each step of the sphingomyelin synthesis pathway, there are multiple paralogs within the *C. elegans* genome that are predicted to perform the same biochemical role (Fig 4a). We investigated independently-derived deletion alleles of the paralogs including the genes we identified in our genetic screen. In every case, only the publicly available deletion allele of the respective mutant gene identified in our screen resisted the effects of wact-190 (Table 1). This suggests that either the specific expression pattern or the biochemical activity of the respective paralog identified in our screen is important for mediating sensitivity to wact-190. Beyond testing the respective deletion alleles, we did not investigate *sptl-2* and *ttm-5* further. The *sms-5* null mutant resists wact-190’s crystal formation (Fig 4b-4d), wact-190-associated developmental arrest/lethality (Fig 4d), and the accumulation of wact-190 in the worm as revealed by mass-spectrometry analysis (Fig 4e). The *sms-5* null mutant also resists Evans blue dye penetration into the anterior pharynx when co-incubated with wact-190 (Fig 2e).

We next investigated the tissues in which *sms-5* plays a role in small molecule accumulation. We used a fosmid-based reporter in which SMS-5 is tagged with YFP on its C-terminus (pPRHM1051). We observed SMS-5::YFP to be expressed in only two tissues; the spermatheca and the pharynx. In the pharynx, SMS-5::YFP is expressed in the muscle and marginal cells, with an enrichment at cell-cell junctions, and a clear enrichment in the anterior half (corpus) relative to the posterior half (isthmus and terminal bulb) (Supplemental Figure 4). Animals mosaic for the transgene show SMS-5::YFP expression in the marginal cells (Fig 4f and 4g).

Wild type animals accumulate wact-190 crystals specifically in the anterior marginal cells. We therefore tested whether expression of SMS-5 specifically in the marginal cells was sufficient to rescue the mutant’s resistance to wact-190 crystal formation. We used a promoter to express an SMS-5::FLAG::mCherry fusion protein (pPRZH1138) specifically in the marginal cells of the anterior pharynx (see Supplemental Figure 5 for details). We found that expression of SMS-5 in the anterior marginal cells can rescue the *sms-5* null phenotypes, including its resistance to both wact-190’s lethality and crystal formation (Fig 4d). Together, these data indicate that SMS-5 functions in the anterior pharynx to make the marginal cells especially sensitive to the accumulation of select small molecules.

**Figure 5.**
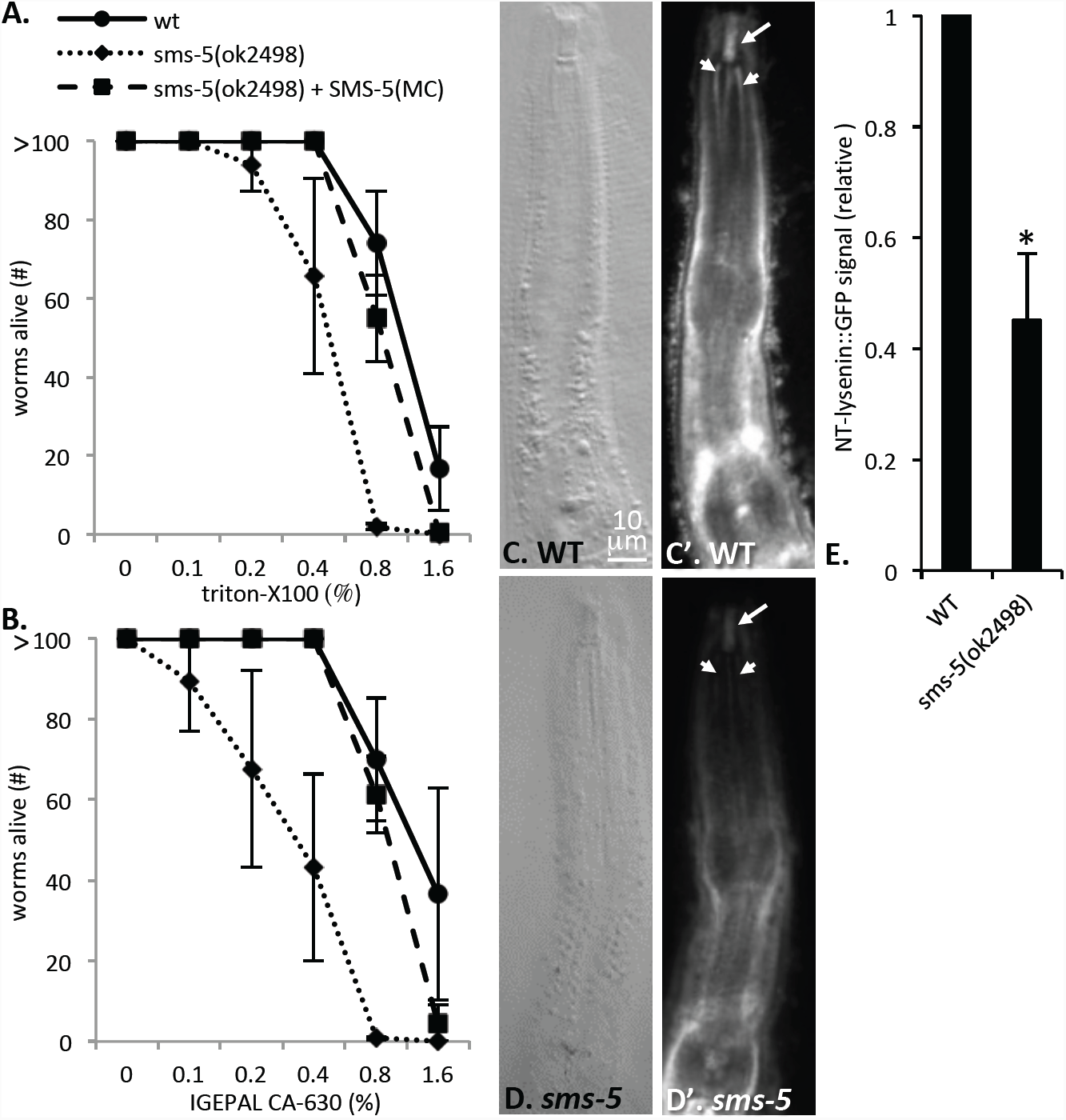
*sms-5* Mutants have Decreased Sphingomyelin Abundance in the Anterior Pharynx. **A-B.** The *sms-5* deletion allele is hypersensitive to the lethality induced by detergents triton-X 100 and IGEPAL CA-630, shown as a v/v percentage of the liquid media. In both cases, differences are significant (*p*<0.05) at a detergent concentration of 0.8% (w/v). SMS-5(MC) represents marginal-cell specific expression of SMS-5 from the *trIs104* integrated transgenic array. **C-D.** Wild type and *sms-5(ok2498)* mutant animals are fixed and stained with the GFP-NT-lysenin protein probe that recognizes clustered sphingomyelin. White arrows indicate buccal cavity and white arrowheads indicate anterior channels. The scale is the same for all panels. **E.** Quantification of the relative GFP signal from more than five biological replicates of GFP-NT-lysenin staining (see methods for details). Asterisk (*) indicates a significant difference (*p*=0.024). Standard error of the mean is shown.

### SMS-5 Contributes to Sphingomyelin Abundance in the Anterior Pharynx

Sphingomyelin facilitates a dense packing of lipids within a membrane [22]. Consequently, sphingomyelin abundance is correlated to a resistance to solubilization by detergents [23, 24]. We therefore tested whether *sms-5* mutants are more sensitive to detergents compared to wild-type animals. As predicted, we observed that the *sms-5* mutant is hypersensitive to detergents compared to wild type controls. Furthermore, expression of SMS-5 from the anterior marginal cells was sufficient to rescue this hypersensitivity to the detergents (Fig 5a and 5b).

We next investigated the relative abundance of sphingomyelin in the anterior pharynx using a protein probe called GFP-NT-lysenin that binds to clustered sphingomyelin (see methods; Supplemental Fig 6) [25, 26]. In wild type fixed worms, we observed an enrichment of GFP-NT-lysenin in tissues surrounding the pharynx, in the buccal cavity and in the channels of the anterior marginal cells (Fig 5c). The *sms-5* null mutant shows a significant decrease of fluorescent signal in the anterior pharynx (Fig 5d and 5e). We infer that the remaining signal is either background staining from the sphingomyelin probe and/or sphingomyelin that is produced by other sphingomyelin synthases [27]. We conclude that *sms-5* mutants produce less sphingomyelin in the anterior pharynx compared to wild type animals.

**Figure 6.**
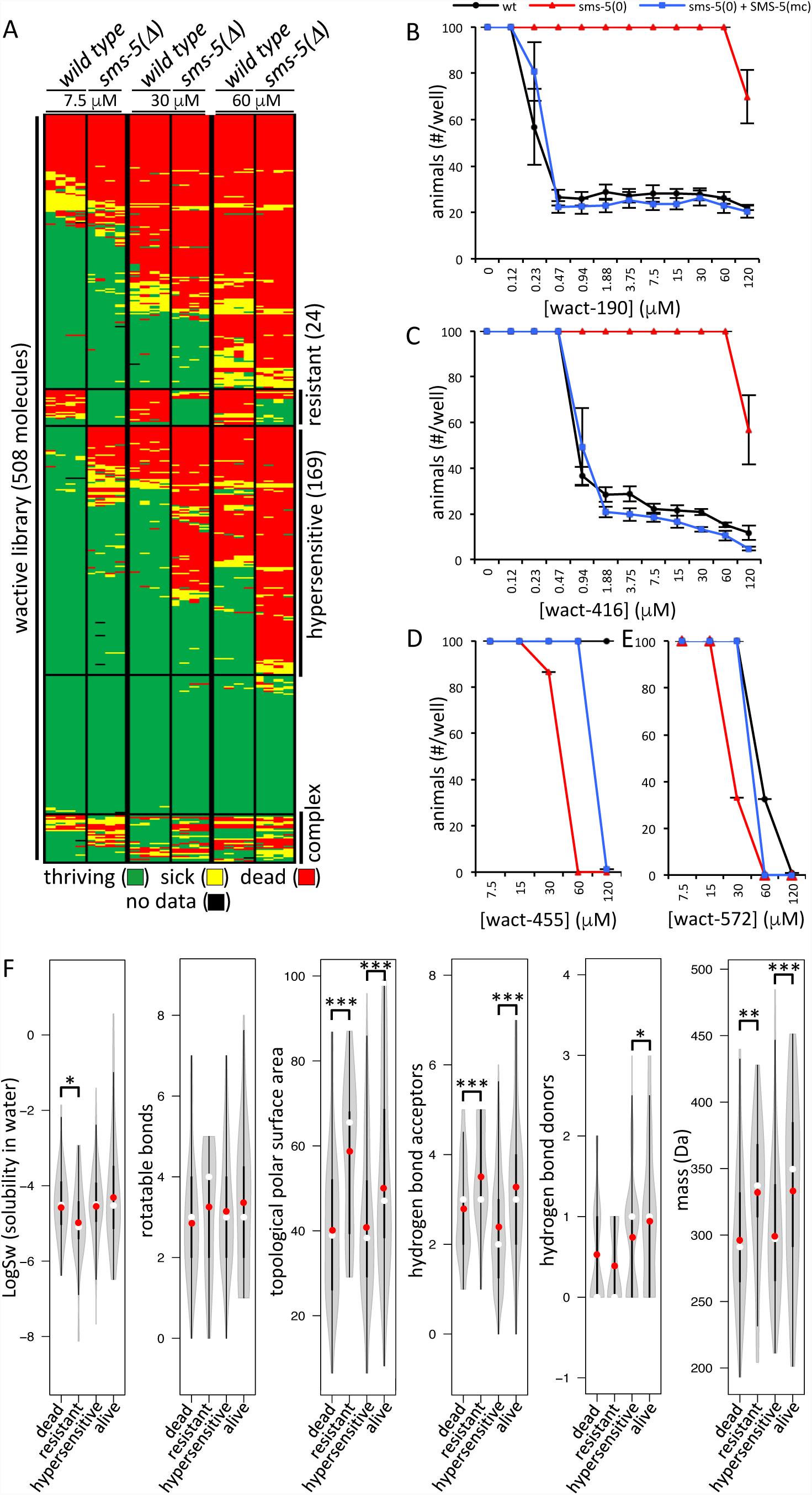
*sms-5* Mutants have Altered Sensitivity to Small Molecules. **A.** A systematic chemical-genetic interaction analysis. Survival assays of animals of the indicated genotype (top) grown in liquid culture in quadruplicate in 508 different wactive small molecules at either 7.5, 30, or 60 μM. Assays were performed in 96-well plates and each well was seeded with 20 synchronized first stage larval hatchlings. Wells were inspected after six days of growth; wells that had 50 or more animals are shown in green, wells that have between 11 and 50 animals are shown in yellow and wells with 11 or fewer animals are shown in red. The resulting population growth of each of the four replicates is shown in each column. Each row corresponds to a distinct wactive molecule. The data is clustered along the y axis. The 24 wactives with reduced potency in the mutants we refer to as ‘resistant’ wactives, and the 169 wactives with increased potency in the mutants we refer to as ‘hypersensitive’ wactives. These are indicated on the right of the chart. **B-E.** Dose-response analyses of *C. elegans* population growth with two molecules of the ‘resistant’ class (wact-190 and wact-416), and two molecules of the ‘hypersensitive’ class (wact-455 and wact-572). Experiments were performed in quadruplicate in 96 well-plate format with ∼50 synchronized L1s deposited into each well (see methods). Standard error of the mean is shown. The *sms-5(ok2498)* deletion mutant (‘*sms-5(0)*’) was used in this analysis, along with the rescuing *sms-5* transgene whereby fluorescently-tagged SMS-5 is expressed exclusively from the anterior marginal cell-specific promoter from the *trIs104* integrated transgene. Resulting wells with fewer than 50 animals in the wact-190 and wact-416 conditions were invariably arrested or dead L1s. **F.** Violin plots comparing the physical-chemical properties of molecules that are classified as either lethal in all conditions (dead), ‘resistant’ (see above), ‘hypersensitive’ (see above), or not obviously bioactive (alive) at 30 μM. See the legend of Figure 1 for a detailed description of the violin plot. One, two and three asterisks indicate a significant difference (*p*<0.05)(*p*<0.01)(*p*<0.001), respectively, between the indicated datasets.

### SMS-5 Expression in the Anterior Marginal Cells Promotes Access to Large Polar Hydrophobic Molecules but Limits Access to Relatively Smaller Molecules

We investigated whether *sms-5* mutant animals are specifically resistant to wact-190, or whether the mutant exhibits resistance to structurally diverse small molecules. To do this, we examined the viability of the wild type and the *sms-5(ok2498)* null mutant when incubated in 508 wactives at three different concentrations (7.5, 30 and 60 μM). In addition to wact-190, we found that the *sms-5* mutant is resistant to 23 other wactives in at least one of the concentrations tested (Fig 6a). We refer to these 24 molecules as ‘resistant’ wactives, the majority of which are structurally distinct. Detailed analyses with two resistant wactives show that SMS-5 expression specifically in the anterior marginal cells rescues this resistance (Fig 6b and 6c).

We were surprised to find that at least 169 (33%) of the wactives killed the mutants at a concentration that had little-to-no effect on wild type (Fig 6a). Because the *sms-5* mutants are hypersensitive to these 169 molecules, we refer to these as ‘hypersensitive’ molecules. Detailed analyses with two ‘hypersensitive’ molecules show that SMS-5 expression specifically in the anterior marginal cells can rescue hypersensitivity (Fig 6d and 6e).

We investigated the physical-chemical properties of the resistant and hypersensitive wactives. We found that the ‘resistant’ molecules have a larger mass, have a greater topological polar surface area (PSA), are significantly less soluble in water, and have more hydrogen bond acceptors, than the molecules that kill both the wild type and the *sms-5* mutant (*p*<0.05) (Fig 6f). The hypersensitive molecules have nearly the opposite set of properties; they are of lower molecular weight, have a smaller PSA, and have less hydrogen bond acceptors and donors than control molecules that fail to kill either the wild type or the *sms-5* mutant (p<0.05) (Fig 6f). We infer that SMS-5 contributes to a marginal cell barrier that normally allows the accumulation of large hydrophobic molecules, but limits the accumulation of relatively smaller and less hydrophobic molecules.

### Some Crystallizing Wactives May Kill *C. elegans* through Multiple Independent Mechanisms

Several lines of evidence are consistent with the idea that wact-190 crystals induced developmental arrest and/or lethality. First, the dose at which crystals form is coincident with the dose that kills worms (Fig 2a). Second, the plasma membrane is disrupted in the same cells in which crystals form (Figs 1 and 2). Third, *sms-5* mutants coincidentally resist the crystal formation and lethality induced by wact-190 (Figs 4d, 6b, and 6c). Fourth, the restoration of SMS-5 activity specifically in the cells in which crystals normally form restores both crystal formation and lethality to the *sms-5* mutant (Figs 4d, 6b, and 6c).

We therefore expected that all of the crystal-forming compounds would behave like wact-190. That is, we expected that all crystal-forming compounds would: i) fail to form crystals in the *sms-5* mutant, and coincidentally, ii) fail to kill the *sms-5* mutant. However, we were surprised to find that 53% of the crystallizing wactives that kill the wild type remain effective at killing the *sms-5* mutant (Fig 7a).

**Figure 7.**
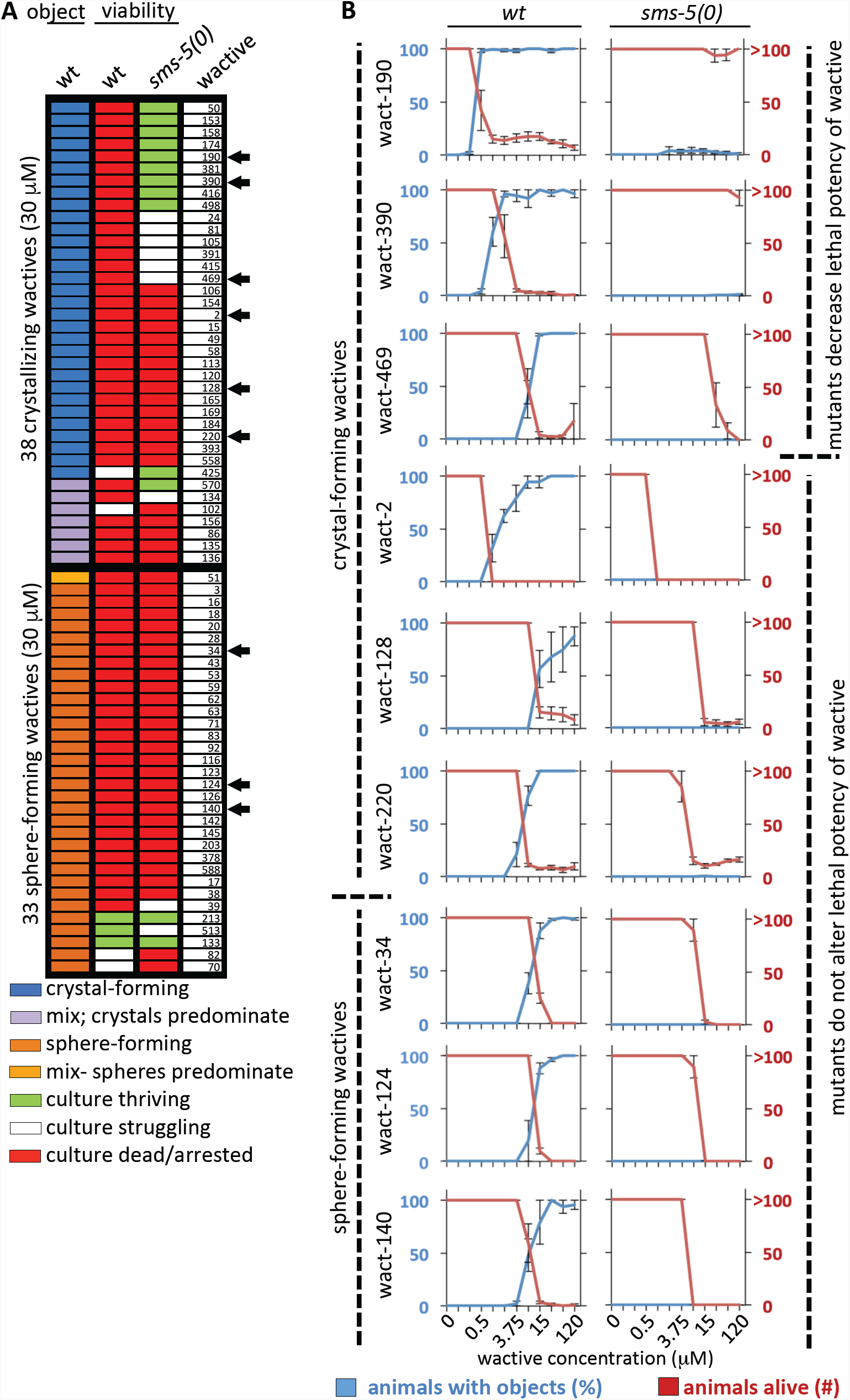
A Comparison of the Lethality and Object-Forming Capability of Wactives in Animals Lacking SMS-5. **A.** The behavior of molecules that form objects in at least 25% of the population are analyzed with respect to their lethal potency in wild type and the *sms-5(ok2498)* deletion mutant. The wactive number is shown on the right and black arrowheads indicate the wactives analyzed in ‘B’. **B.** Dose-response analyses for select crystal and sphere-forming wactives in the indicated genetic background (indicated at the top of the two columns of graphs). Each of the wactives is analyzed over a two-fold dilution series of concentrations (indicated at the bottom of the three columns of graphs). Wactive molecules are indicated on the left. The standard error of the mean is shown. See methods for more details.

To investigate the discordance of the ability to kill the *sms-5* mutant among crystal-forming wactives, we investigated a handful of crystal-forming compounds in detail for their ability to kill and form crystals in the *sms-5* mutant. In all cases, we found that the *sms-5* mutant resists crystal formation, but half of these wactives are still able to kill the mutant (Fig 7b). This suggests that while some crystal-forming wactives may kill only via crystal formation, others may kill via two mechanisms-one that is crystal-dependent and another that is crystal-independent.

We also performed similar dose-response analyses for a handful of sphere-forming compounds. In all cases examined, *sms-5* mutants remained sensitive to these wactives despite a lack of sphere formation (Fig 7b). This suggests that the sphere formation itself is simply a visible marker of compound accumulation and is not directly related to how the sphere-forming wactives kill the animals.

### *sms-5* Mutants are Hypersensitive to the Loss of Cholesterol

*C. elegans* is a cholesterol auxotroph, requiring as little as 2.5 ng/ml to survive [4]. CHUP-1 (initially called CUP-1) is a key transmembrane protein involved in cholesterol absorption in *C. elegans* [28]. Consequently, *chup-1* mutants are hypersensitive to cholesterol-limited conditions [28]. CHUP-1 is expressed in the posterior pharynx but is more prominently expressed in the intestine [28]. Despite *C. elegans*’ reliance on mature sterols for development [4], *chup-1* null mutants are viable, suggesting that additional mechanisms of cholesterol uptake exist [28].

Given SMS-5’s role in the accumulation of hydrophobic small molecules, we asked whether SMS-5 might function in parallel with CHUP-1 to help absorb cholesterol, which is also a small hydrophobic molecule. Wild type hatchling parental (P0) animals placed on cholesterol-limited plates are themselves able to grow to adulthood, likely because of maternally-contributed cholesterol stores [29]. However, these P0 animals produce fewer F1 progeny and these F1s are slow to develop to the L4 stage (see Fig 8a, first column). We tested how the *sms-5* null mutant behaves on cholesterol-limited plates. Like the *chup-1* null control, *sms-5* null animals had difficulty growing on cholesterol-limited conditions (Fig 8a; Supplemental Fig 7). The growth deficit of the *sms-5* mutant is rescued by the expression of SMS-5 specifically in the anterior marginal cells (Fig 8a; Supplemental Fig 7). Furthermore, the *chup-1; sms-5* double mutant grows worse on cholesterol-limited conditions than either single mutant (*p*<0.05) (Fig 8a; Supplemental Fig 7). This double mutant analysis indicates that SMS-5 may function in parallel to CHUP-1 in the absorption of cholesterol.

**Figure 8.**
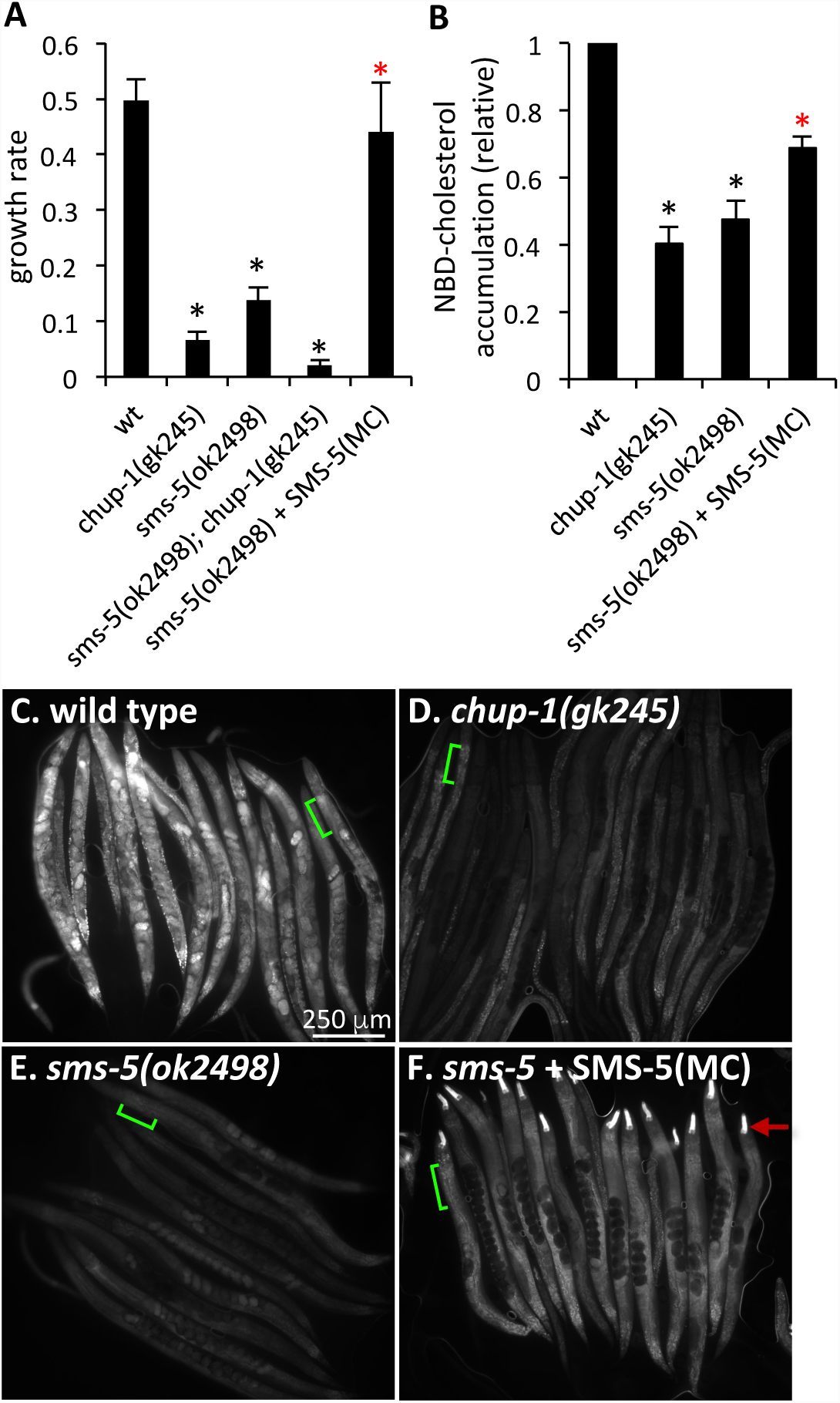
SMS-5 is Required for Development in Cholesterol-Limited Conditions. **A.** The growth rate of animals of the indicated genotype grown on cholesterol(xol)-limited plates (containing 50 ng/mL xol) is reported relative to the same strains grown on plates with standard concentrations of cholesterol (5000ng/mL xol) (y-axis). See methods for additional details. Asterisks indicate significance (*p*<0.001) relative to the wild type control; the red asterisks indicate significance (*p*<0.05) relative to the *sms-5(ok2498)* deletion mutant. The double mutant has a more severe growth defect relative to either single mutant (*p*=0.06). **B.** Accumulation of NBD-cholesterol signal of the indicated mutants relative to wild type controls. Black asterisks indicate significance (*p*<0.01) relative to the wild type control; the red asterisks indicate significance (*p*<0.05) relative to the *sms-5(ok2498)* deletion mutant. In both A and B, standard error of the mean is shown. **C-F**. Images of multiple animals after the indicated strains were incubated in NBD-cholesterol for six days. The scale is the same for all images. Green lines exemplify the area used in each animal to calculate signal in B. Red arrow indicates the expression of YFP that is restricted to the anterior pharynx that is used as a marker to follow the rescuing transgene *trIs104*. In A, B, and F, SMS-5(MC) represents marginal-cell specific expression of SMS-5 from the *trIs104* integrated transgenic array. Supplemental Figure 8 shows that in the absence of NBD-cholesterol, each strain auto-fluoresces in the GFP channel to the same extent.

We also investigated whether the *sms-5* mutant is deficient in cholesterol absorption by examining the uptake of a fluorescent analog called NBD-cholesterol, which was previously used to characterize *chup-1’*s role in cholesterol uptake [28]. Similar to previous results, we observed an obvious decrease in NBD-cholesterol accumulation in the *chup-1* mutants relative to wild type control after synchronized L1s were incubated in NBD-cholesterol for six days (see methods) (Fig 8b-8d). We observed a similar decrease in NBD-cholesterol accumulation in the *sms-5* mutant (Fig 8e), which could be rescued by expressing SMS-5 specifically in the anterior marginal cells (Fig 8f). Together, the data indicate that SMS-5 contributes to cholesterol absorption.

## Discussion

Here, we show that the marginal cells within the anterior pharynx have distinct properties that facilitate unique interactions with exogenous small molecules. We can speculate why these marginal cells play a special role in the animal by first considering that nematodes are cholesterol auxotrophs and that free cholesterol is relatively rare within the environment [1, 4]. The marginal cells form the channels through which excess ingested fluids are expelled from the animal [7, 9]. Hence, the marginal cells may function as a nutrient salvage system to ensure that precious small molecules are not again lost to the environment. This model is consistent with several observations: First, exogenous small molecules visibly accumulate almost exclusively in the cells that line the anterior channels. Second, the pharynx is the first tissue to accumulate a fluorescent analog of cholesterol in time-course analyses [30]. Third, the anterior pharynx accumulates obviously higher levels of cholesterol compared to the posterior pharynx [10]. Finally, absorption of these small molecules is SMS-5-dependent and sphingomyelin-rich membrane has been repeatedly shown to act as a sink for cholesterol [24].

Exactly how SMS-5 facilitates the absorption of hydrophobic molecules is not clear. We do know that sphingomyelin incorporation into a lipid bilayer alters the physicochemical properties of that bilayer (reviewed in [31]) [32]. In other animals, SM is tightly associated with cholesterol within the plasma membrane (PM) and creates sub-domains of densely packed regions within the lipid bilayer [32]. Whether or not SM physically associates with mature sterols or metabolites thereof within the PM of nematodes remains an open question [4]. Given that cholesterol and other mature sterols are very hydrophobic, PM that is enriched for SM-associated sterols could act as a sink for other hydrophobic molecules. By contrast, a reduction in sphingomyelin abundance decreases the packing density of lipids [23, 24] and likely makes it easier for lower molecular weight molecules to pass through the PM barrier. Hence, the biological properties of SM are consistent with our findings that the absence of SMS-5 hinders the accumulation of larger hydrophobic wactives while simultaneously facilitating an increase in potency of smaller wactives.

Our discovery that small molecules may kill worms through crystallization has implications for drug screens using *C. elegans* as a model system. As a group, crystallizing wactives have structural variety and varied potencies, but all have the potential to kill or arrest young larvae through either perforation of the plasma membrane and in some cases, blockage of the alimentary track. For many of these crystallizing wactives, both crystallization and lethality is dependent on an intact SMS-5 sphingomyelin synthesis pathway. For simplicity’s sake, we refer to these molecules as ‘class I’ crystallizing wactives. Class I crystallizing wactives may have no mechanism of action aside from crystallization. Our unpublished observations suggest that class I crystallizing wactives do not have wide-ranging utility against diverse parasitic nematodes. Hence, our future screens for novel nematicides using *C. elegans* as a model system will include a step to determine whether the wactive’s bioactivity is SMS-5-dependent.

There is a subset of crystallizing wactives whose lethality is not dependent on SMS-5. Again for simplicity’s sake, we refer to these molecules as ‘class II’ crystallizing wactives. In an *sms-5* mutant background, the crystallization of class II wactives is abolished, but their lethality persists. This suggests that class II crystallizing wactives may have at least two mechanisms of killing-one that is crystal-dependent, and one that is crystal-independent. Knowing that class II wactives exist has important ramifications for how we investigate the mechanism-of-action of bioactive molecules. Typically, our first approach towards characterizing a wactive’s mechanism-of-action is to screen for *C. elegans* mutants that resist the molecule’s lethal effects. When successful, the mutant gene that confers resistance provides insight into the molecule’s mechanism of action [12-14, 33]. However, with wactives that have the potential to kill via crystal-dependent and crystal-independent mechanisms, any worm with a mutation that confers resistance to one mechanism will likely be killed by the second mechanism. Isolating a mutant that resists both mechanisms of action would be rare, occurring roughly once in every four million mutant genomes if loss-of-function mutations were sufficient to confer resistance to each lethal mechanism (and even rarer if resistance is conferred by gain-of-function only). Knowing that a bioactive molecule behaves as a class II crystallizing wactive may facilitate the characterization of its mechanism of action by screening for resistant mutants in an *sms-5* null background wherein crystal formation is abolished.

Investigating the sensitivity of the *sms-5* mutant to the entire wactive library revealed that this mutant is at least two-fold more sensitive than wild type to more than a third of the small molecule structures surveyed. Detailed analysis with a small subset of these ‘hypersensitive’ molecules indicates that restoring SMS-5 function to only the marginal cells is sufficient to confer normal sensitivity to these wactives. These observations raise several important points. First, it raises the possibility that the tissue of the anterior pharynx is either the target of many bioactive molecules and/or serves as the conduit through which many bioactive molecules traverse en route to their targets in other tissues. Intriguingly, there are several sphere-generating wactives that fail to generate spheres in the mutant background, but simultaneously increase in potency in the mutant background. This suggests that without SMS-5 in the pharynx, the sphere-generating wactives cannot be contained within the pharynx and are better able to disrupt targets throughout the animal. Second, the *sms-5* mutant becomes a tool with which we can increase the potency of many small molecules. Given that the *sms-5* deletion mutant is otherwise healthy, it is a useful sensitized background for future small molecule screens and other chemical genetic experiments.

## Acknowledgements

We thank Greg Fairn for reagents and guidance in characterizing sphingomyelin abundance and localization in the worm. We thank Kevin Chan for performing some microinjections and integrating some transgenes; David Hall, Zeynep Altun, and Chris Crocker for permission to modify Worm Atlas schematics; Andrew Burns for helpful comments on the work; Andy Fraser and Michael Shapiro for whole genome sequence analysis. We thank Lindy Holden-Dye and Fernando Calahorro Nunez for preliminary analyses. We thank the *C. elegans Genetics Centre* and Shohei Mitani for mutant strains. This work is supported by CIHR grants (376634 and 313296) and a CRC to PJR, and an NIH grant (OD 010943) to DHH.

## Materials and Methods

### Crystal and Sphere Analyses

Synchronized L1s were added to wells of a 96 well-plate (50 L1s / well) containing liquid NGM, *E. coli* (HB101) as food source (OD600 = 1.2) and 30 μM of 240 wactives in duplicate wells. 48 hours later, worms were transferred to Eppendorf tubes, and washed once with fresh M9 and spun down at 3000 rpm for 1 minute to form a tight worm pellet of ∼10 μL. Then, 5 μL of 50 mM levamisole (to a final concentration of ∼ 16.7 mM) was added to paralyze worms. Worms were then mounted on a 2% agarose pad on a glass slide. Live worms were then observed for presence of birefringent crystals or non-birefringent spheres in the pharynx using 40 X objective of a Leica DMRA microscope. For both the survey of 238 molecules and the dose response analyses, a minimum of three biological replicates were processed, and at least 20 worms per replicate were counted for the presence or absence of spheres and/or crystals.

### *C. elegans* liquid-based 6-day viability assays

The 6-day viability assay methods were adapted from a previously described liquid-based chemical screening protocol [14]. Briefly, a saturated culture of HB101 *E. coli* was concentrated 2-fold in liquid Nematode Growth Medium (NGM, Burns et al., 2015 for recipe) and 40 μL of bacterial suspension was dispensed into each well of a flat-bottom 96-well plate. Compounds and DMSO controls were pinned into the 96-well plates using a 96 pin replicator with a 300 nL slot volume (V&P Scientific). The final concentration of DMSO in each well was 0.6% v/v. Approximately 20 synchronized L1 larval stage worms were added to each well of the 96-well plates in 10 μL of M9 buffer (see [34] for recipe). For some experiments, approximately 50 synchronized L1 stage worms were added to 80 μL of bacterial suspension in 20 μL of M9 buffer. Worms were synchronized to L1 stage through hypochlorite bleaching of gravid adults performed the previous day (Burns et al., 2006 for protocol). Assay plates were sealed with Parafilm and incubated for 6 days at 20 °C with shaking at 200 rpm. After 6 days, the plates were observed using a dissection microscope and the wells were categorized according to the number of viable adult and larval stage worms in each well. Wells with more than 50 animals after 6 days were categorized as over-grown. The number of worms in wells with approximately 50 or fewer worms were counted. All viability assays were completed in quadruplicate.

To analyze the response of various strains to detergents, 25 synchronized L1-larvae were added to each well of a 24-well plate seeded with 25 μL OP50 and containing a final concentration of 0-1.6% detergent (Triton-X-100 or IGEPAL-630) dissolved in water (day 0). On day 6, the number of worms alive was counted and recorded. Results are from three independent trials with four technical replicates.

### Analysis of Evans Blue Dye penetration into the pharynx

L1-stage worms were incubated with 1% DMSO (control) or 60 µM wact-190 (in 1% DMSO) in liquid for 24hrs. Worms were washed once with M9 and incubated with a final concentration of 0.1% Evans Blue Dye in 500 µL of M9 in siliconized Eppendorf tubes for 4 hours. Worms were then washed three times with M9 solution, suspended in 10 µL M9 and paralyzed for microscopy by adding 4 µL of 50 mM levamisole. Live worms were mounted on 3% agarose pad. Dye fluorescence was observed in TX2 channel of a Leica DMRA microscope at 630X total magnification and quantified using Fiji (ImageJ) software by calculating the Mean Gray Value in the anterior pharynx relative to the Mean Gray Value in a nearby space in the frame devoid of worms.

### Molecular Biology

**pPRHM1051** is the wrm0626dC03 fosmid, which contains the *sms-5* locus (W07E6.3), wherein the SMS-5 coding sequence is C-terminally tagged with YFP. This construct rescues mutant *sms-5*’s resistance to wact-190. pPRHM1051 was created using Oliver Hobert’s fosmid tagging methodology [35]. **pPRZH1138** (pgp-14p::SMS-5B(genomic)::FLAG::mCherry). This construct is based upon pPRHM1065 (myo-2p::SMS-5::FLAG::mCherry). The *pgp-14* promoter was amplified by PCR from pPRZH1160 and used to replace the *myo-2* promoter from pPRHM1065. pPRZH1138 rescues mutant *sms-5*’s resistance to wact-190. Detailed construct plans are available upon request.

### Mass Spectrometric Analyses of Wact-190 Accumulation

#### Worm preparation

We grew a mixed stage population of wild type (N2) *C. elegans* and performed and egg-prep as previously described [34]. Embryos were allowed to hatch overnight in M9 buffer. The resulting synchronized L1s were harvested the next morning. L1s were then grown on 10 cm plates, with 10,000 L1s per plate, and incubated at room temperature for 48 hours. Worms were then washed off the two plates with M9 buffer, collected into a 15 ml conical tube and washed three times. In parallel to this, we prepared a ‘drug’-incubation buffer by first inoculating 400 ml of LB with 50 μl of an overnight culture of E. coli (HB101), grew it overnight, and centrifuged 50 ml of this fresh culture for 10 min at 3200 rpm. We decanted the LB and rinsed the bacteria in ∼50 ml of NGM buffer once and then resuspended the bacteria in 25 mls of NGM. We resuspended 20,000 worms in 1 ml of the NGM + HB101 solution in either 1% DMSO or 30 µM of small molecule (to a final DMSO concentration of 1%) in 1.5 ml siliconized microcentrifuge tubes. We then incubated the tubes for 4 hours at 20°C on a nutator. Thereafter, we washed the worms five times with ice-cold M9 buffer, keeping the samples on ice as much as possible. We removed the M9 and flash froze the samples using liquid nitrogen and stored the worms at −80°C until the samples were ready to process by mass spectrometry.

#### MTBE Extraction

We first lyophilized the worm pellet and resuspended it in 200 μL of 0.1M NaCl, followed by sonication for 5 min at 100W in a Misonix cup sonicator. Samples were then spiked with 43 ng of internal standard (triamcinolone acetonide) and extracted twice with methyl tert-butyl ether (sample: MTBE, 1:5, v/v). Organic layers were pooled and evaporated to dryness under N_2_. Extracts were re-suspended in 200 µL of HPLC-grade methanol and analyzed as described below.

#### LC/MS/MS Analysis

The samples were analyzed by LC/MS/MS using a 6410 LC/MS/MS instrument (Agilent Technologies) with an ESI source in positive ion mode. Samples were separated on a Zorbax XDB-C18 column (4.6 × 50 mm, 3.5 µm) at 0.4 mL/min. The mobile phase consisted of HPLC-grade water (A) and methanol (B) both containing 5mM NH_4_Ac. The following gradient was run: 0-1 min, 60% (B); 1-3.3 min, 60% to 100% (B); 3.3 – 7 min, 100% (B); stop time, 10 min; post-time, 5.5 min. MS parameters were as follows: nebulizer pressure 35 psi, drying gas (nitrogen) 10 L/min, VCap 6000 V, Delta EMV 800V, column temperature 40°C and drying gas temperature 350°C for all compounds. Using multiple reaction monitoring the following transitions were measured:

| Compound | RT (min) | MRM | Fragmentor (V) | Collision Energy (V) |
| --- | --- | --- | --- | --- |
| Trimacrinolone Acetonide | 5.7 | 435→415 | 108 | 5 |
| wact-190 | 7.7 | 330→107 | 155 | 22 |

### GFP-NT-Lysenin Protein Production and Staining

GFP-NT-Lysenin protein [36] was produced as previously described [37]. Briefly, BL21(DE3) Rosetta cells were transformed withpMAL-C2-GFP-NT-Lysenin vector (a gift from Dr. Gregory Fairn, University of Toronto) and cultured in 1 L of LB broth at 37°C until OD_600_ reached 0.6. Expression was induced with 1 mM IPTG for 6hrs at 25°C. Cells were frozen overnight at −80°C. Cells were lysed using B-PER reagent (Thermo Fisher Scientific) according to manufacturer’s instructions, in the presence of excess MgCl_2_, DNase I, lysozyme and protease inhibitor cocktail. Crude cell lysate was passed through a Sepharose column containing 5mL amylose resin (New England Biolabs) and the MBP-tagged GFP-NT-Lysenin fusion protein was eluted with fresh 10 mM maltose in 1X PBS (pH 7.4). All purification steps were carried out at 4°C. SDS-PAGE confirmed the presence of the 90 kDa fusion protein, which was stored at 4°C until use.

Synchronized adults were fixed in 1.5 ml Eppendorf tubes using the Modified Finney-Ruvkun protocol [38]. Briefly, worms were washed with chilled M9 buffer three times, removing as much bacteria as possible. The worm pellet was incubated in ice for 30 minutes then mixed with 2X MRWB (5% methanol) and freshly prepared paraformaldehyde (final concentration-4%), freeze-thawed in an ice-ethanol bath four times, and then incubated on ice for 2hrs. The worm suspension was then reduced and oxidized with 1% beta-mercaptoethanol and 10mM DTT and 0.3% hydrogen peroxide respectively. The following changes were made to the protocol: All washes were done with tris-triton buffer (TTB) lacking triton-X-100; triton-X-100 was also replaced with 1X PBS in Buffers A and B in the final steps. The quality of fixation was tested by actin staining of fixed worms following 2hr incubation at RT with 0.328 nM phalloidin.

To stain worms with GFP-NT-Lysenin, 20 µL of fixed worms were suspended in 0.5 mg/mL of GFP-NT-Lysenin in the final volume of 100 µL of 1X PBS in a 0.6 mL tube. Fixed worms were incubated at 4°C for 4 hours in the dark with constant mixing. Worms were washed two times with 5.5 mL ice-cold 1X PBS. 10 µL of worms were mounted on a glass slide containing 3% agarose and a cover-slip was applied. Animals were imaged in the GFP channel using 20X magnification and identical exposure times for all strains. The experiment was repeated at least three times. Fluorescence intensity in the anterior pharynx was quantified using ImageJ software. For each worm, fluorescence intensity was determined by subtracting the background mean gray value (MGV) from MGV of the procorpus. The Student’s t-test was performed on mean values of the different trials using GraphPad 6.0.

### Cholesterol-Limitation Assay

Worms were grown on MYOB plates seeded with E. coli (OP50). Gravid adults were bleached and the resulting embryos were incubated overnight at 20°C to yield synchronized L1 parents (P0s). The next day, ∼80 synchronized L1 P0s were plated onto modified NGM plates (35 mm) in triplicate. The modified NGM recipe is used to deplete its contents of sterols, and is described in [10]. Briefly, the agar is replaced with agarose (Froggarose) and the peptone is extracted with ether three times and dried in a fumehood overnight. The plates incorporate either standard (5000 ng/mL, “+xol”) or low (50 ng/mL, “-xol”) concentrations of cholesterol. We did not prepare plates without cholesterol because resulting F1s arrest as L1s, regardless of the genotype we tested. All plates are then seeded with 50 µL of an overnight OP50 LB culture.

Three days after seeding plates with L1 P0s, 20 gravid adults are transferred to fresh plates in triplicate and allowed to lay eggs. P0s raised on +xol plates are transferred to new +xol plates; P0s raised on -xol plates are transferred to new -xol plates. After 6 hours of egg-laying, the P0s are removed. We define this as ‘day 0’.

On day 4, the number of F1 L4s and/or adults on the plates are counted. The experiment was repeated at least three independent times (each with three technical replicates) at 20°C. For each strain, we measure the impact of xol limitation by calculating the ratio of F1 L4s/adults on the -xol plates relative to the F1 L4s/adults on the +xol plates (to account for fecundity/growth issues unrelated to xol). Note that we store our filter-sterilized cholesterol stock (in 100% ethanol) in a 50 mL falcon tube sealed with parafilm at 4°C to prevent evaporation.

### NBD-cholesterol Accumulation Assay

NBD-cholesterol accumulation assays are based on the modification of a previously published protocol [28]. An overnight OP50 culture was pelleted and diluted with autoclaved water to which was added 22-NBD-cholesterol dissolved in ethanol (Thermofisher Inc) at a concentration of 200 µM and vortexed vigorously. 175 µL of the OP50 + 22-NBD-cholesterol mixture was added to 6 cm NGM Petri plates containing 5 µg/mL (∼13 µM) cholesterol made on the same day and then dried in a laminar flow hood for 1 hour. The next day (defined as ‘day 0’), 25 synchronized L1-stage larvae were plated onto each of three 6 cm NGM Petri plates for each strain. On day 6, worms were removed from plates and washed three times with M9. After the final wash, 200 µL of worm pellet was added to an Eppendorf tube, to which was added 75 µL of 20% of freshly-prepared paraformaldehyde (PFA) diluted to a final volume of 500 µL with M9 (to final concentration of 3%). Worms were fixed for 30 minutes at room temperature. Next, the worm pellet was washed twice with 1 mL of a 100 mM glycine solution to quench the PFA. A minimal volume of concentrated worm pellet (∼5 µL) was then deposited onto a 3% agarose pad on a glass slide. A small drop of a fluorescent mounting medium (Thermo Scientific™ Richard-Allan Scientific™ Cytoseal 280) was added to the sample before adding a coverslip. For each sample, 30-70 worms were imaged using identical exposure times and 10-20X objectives. ImageJ was used to quantify fluorescence intensity in the anterior and posterior halves of the worm gut, along with background areas next to the anterior and posterior halves to control for differences in background signals. The fluorescence intensity is expressed as Mean Gray Value (MGV). The experiment was performed at least three times. Except where noted, worms were incubated at 20°C.

### Emission spectra of wactives

Emission spectra for select wactives were measured by making a 50 µL solution of each compound at a concentration of 250 µM in double-distilled water. A fluorescence intensity scan was performed in 96-well plate format using an Infinite M200 Pro microplate reader (Tecan Life Sciences) with an excitation wavelength set at 390 nm and the range of emission wavelength measured at 425-700 nm with 25 nm steps. The resulting intensity values were normalized against a solvent blank.

### TEM Analyses

Approximately one thousand synchronized L1-stage worms were added to 60 mm plates containing 60 µM of wact-190 or 1% DMSO vehicle control seeded with OP50 bacteria. 24 hours later, worms were collected and prepared for transmission electron microscopy as previously described [39]. Briefly, live animals were loaded into a metal planchette in a slurry of *E. Coli* and fast frozen under high pressure using a Balt-tec HM 010 freezing device, slowly thawed into osmium fixative in acetone, then infiltrated with plastic resin and thin sectioned for views at high resolution using a Philips CM10 electron microscope.

### Statistics and Graphs

Except where indicated, statistical differences were measured using a two-tailed Students T-test. Violin plots were generated using the online tool BoxPlotR (http://shiny.chemgrid.org/boxplotr/).

## Supplemental Figures

**Figure S1.**
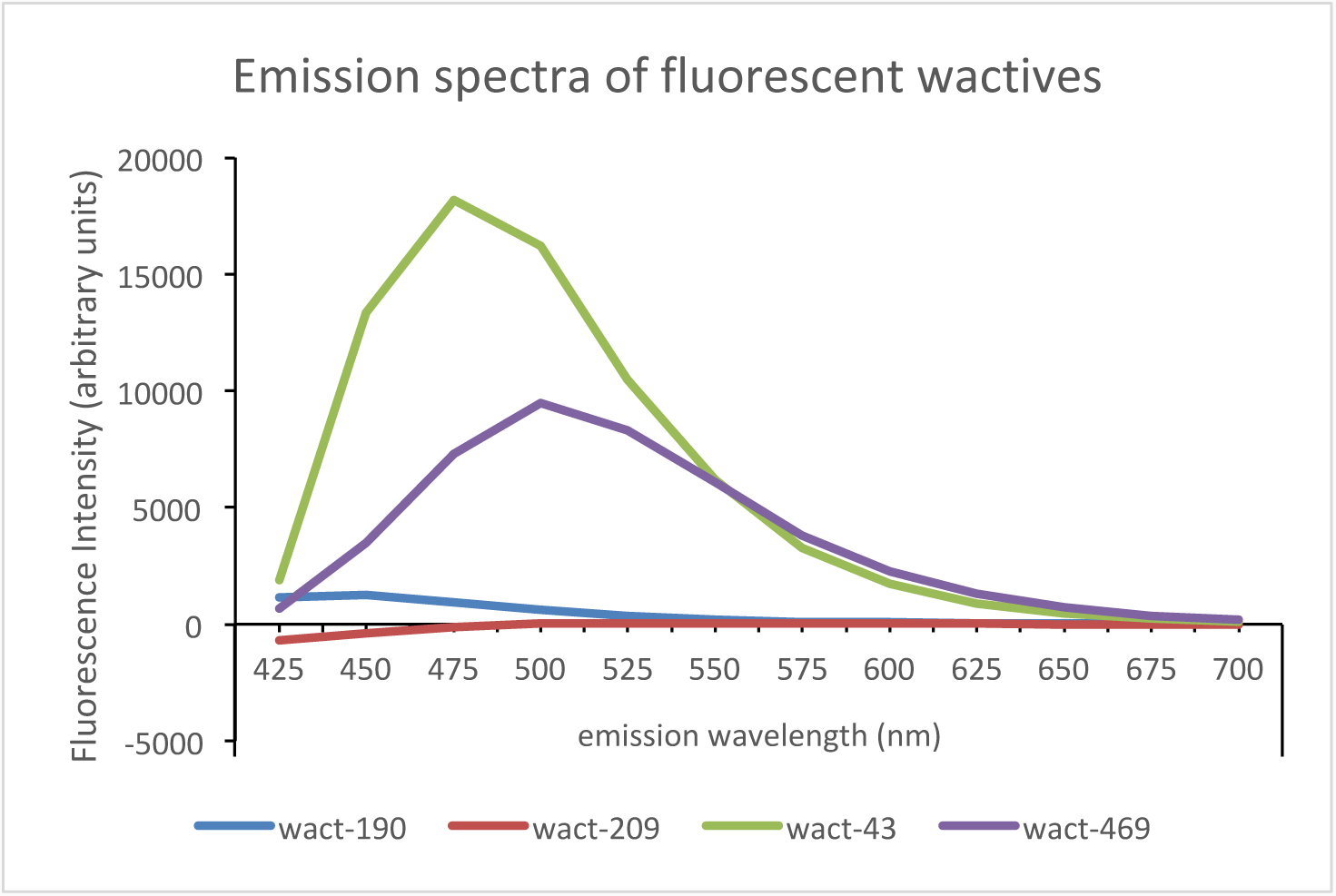
Fluorescent Properties of wact-190, wact-469, wact-209, and wact-43. Wactive compounds were analyzed at a final concentration of 250 μM in 50 μL of double-distilled water in a 96-well flat-bottom plate (Corning). A fluorescence intensity emission scan was performed by exciting the compounds at 390 nm. The resulting emission spectrum was plotted for each compound.

**Figure S2.**
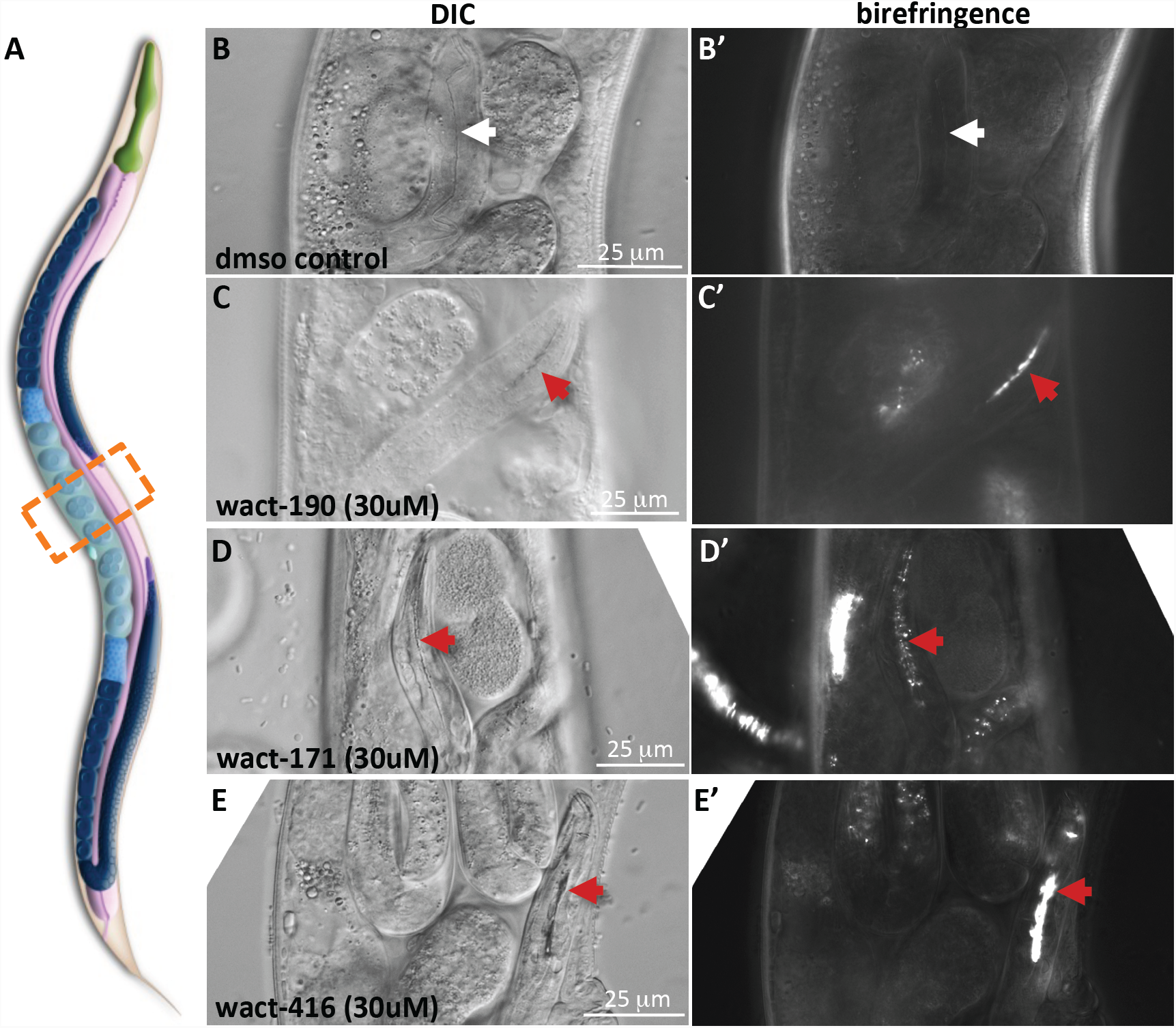
Animals that Hatch Internally can Accumulate Crystals. **A.** A schematic of *C. elegans* (with permission from WormAtlas). The orange box highlights the approximate area shown in figures B-E. **B-E.** Differential interference contrast (DIC) are shown on the left column and the corresponding birefringent images are shown on the right. All animals shown were grown in liquid as synchronized L4s, incubated in either vehicle control (1% DMSO), or the indicated wactives, for 48 hours. The anterior pharynx of the internal hatchlings is indicated with a white arrow (no obvious objects) or a red arrow (where crystals are evident).

**Figure S3.**
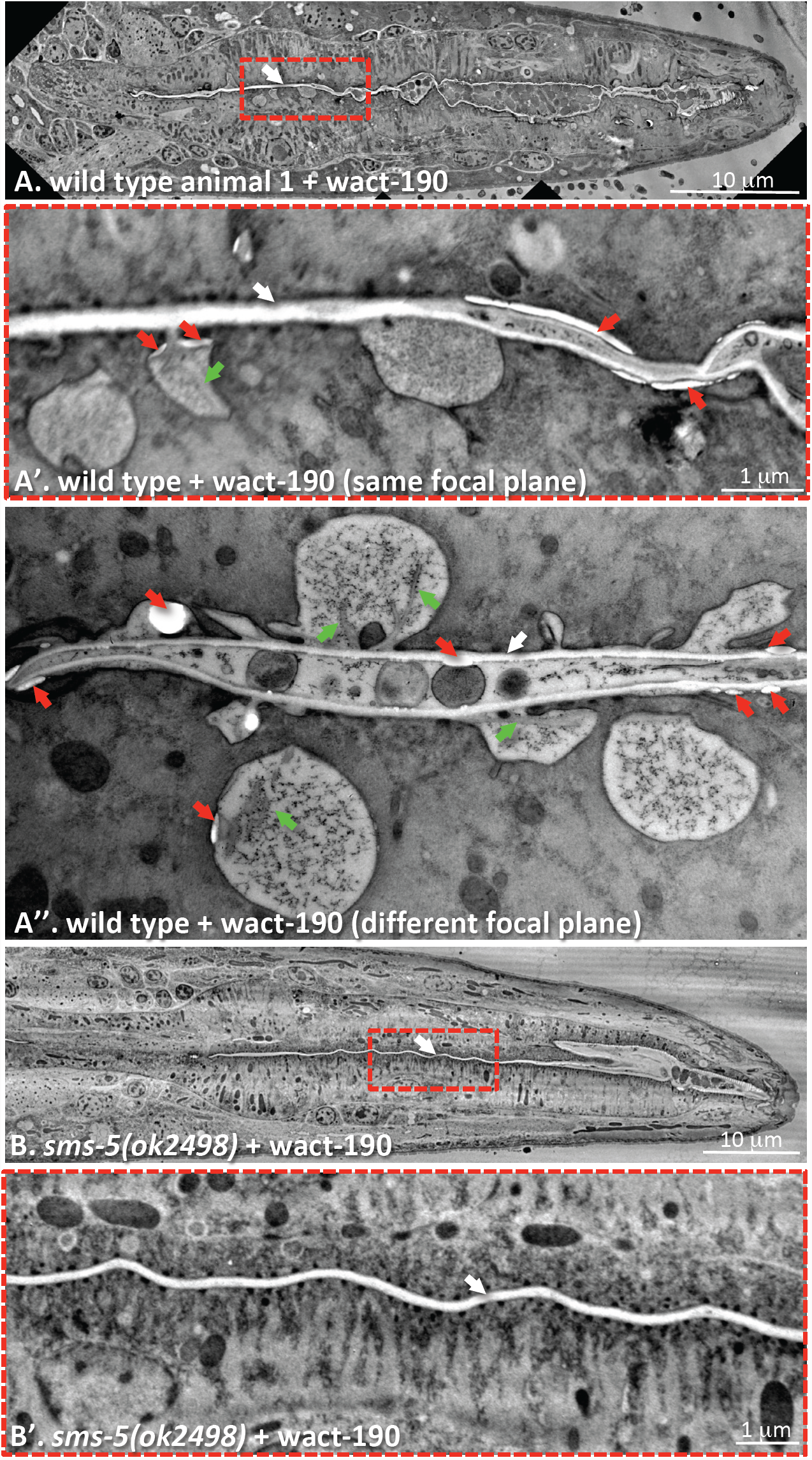
Transverse Transmission Electron Microscopy (TEM) Sections of Animals Co-Incubated with wact-190. **A.** A wild type L1 animal incubated in 60 μM of wact-190 for 24 hours. White arrows indicate apical membrane of the marginal cells, green arrows indicate crystal-like objects, and red arrows indicate unusual space that is consistent with either wact-190 accumulation or tissue separation due to wact-190 treatment. A’ is an enlarged area boxed in ‘A’; A’’ is another focal plane of the pharynx of the same animal. Given that we fail to see wact-190-related spheres *in vivo*, we interpret the circular inclusions in the transverse T.E.M. images to be caused by crystal growth. **B.** An *sms-5(ok2498)* mutant L1 animal incubated in 60 μM of wact-190 for 24 hours and then examined by TEM. B’ is an enlarged area boxed in ‘B’.

**Figure S4.**
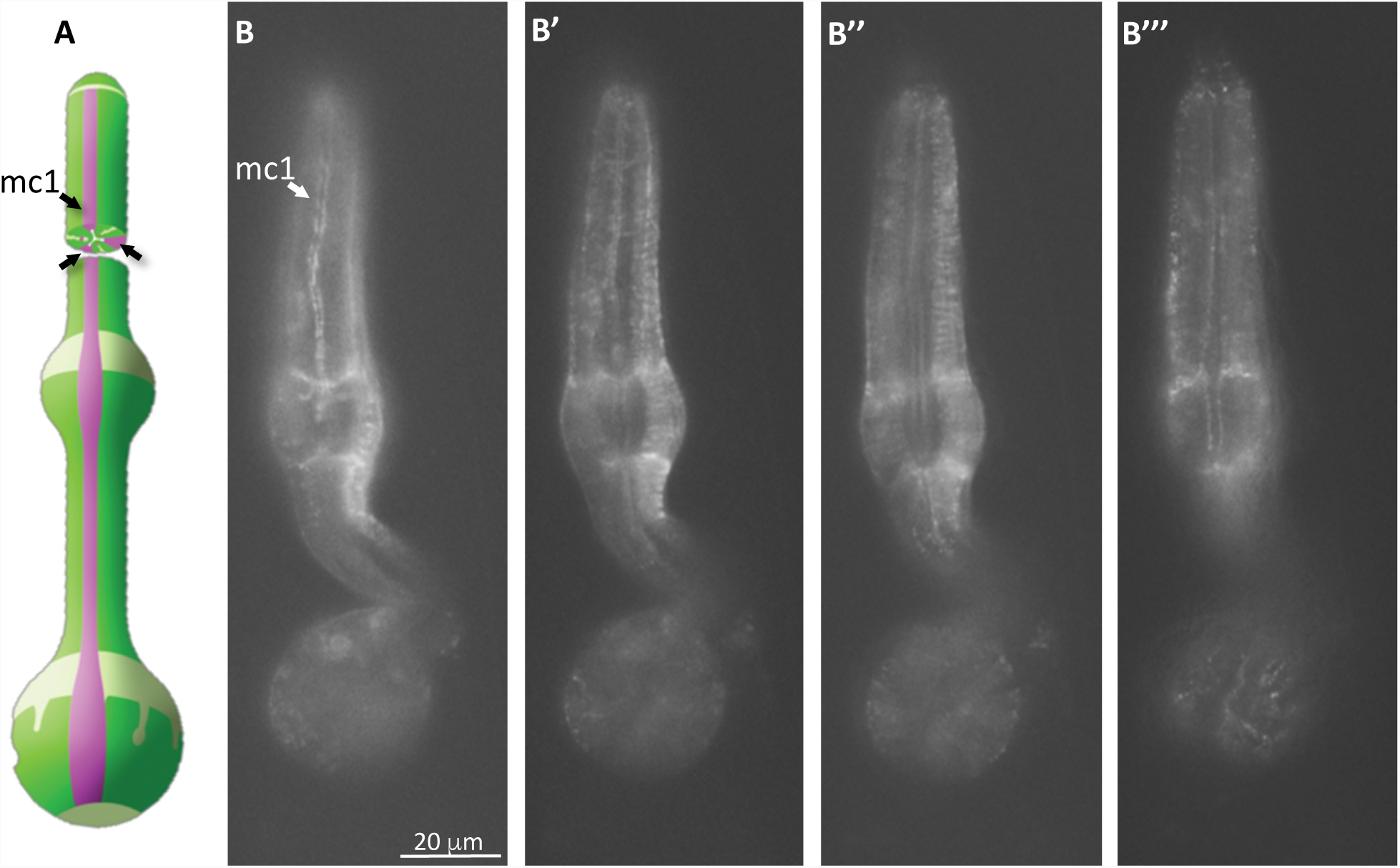
The Expression Pattern of SMS-5 in the Pharynx. Animals harbouring an extrachromosomal array with the pPRHM1051 transgene (which has YFP inserted in frame at the C-terminus of the *sms-5* coding sequence in the context of the WRM0626dC03 fosmid) show fluorescent signal in only the pharynx and spermatheca (not shown). **A.** A schematic of the pharynx (courtesy of WormAtlas) indicating the marginal cells (pink) with black arrowheads. **B.** Four different focal planes of the pharynx of a non-mosaic animal expressing the SMS-5::YFP transgene. A marginal cell (mc1) is indicated with a white arrowhead.

**Figure S5.**
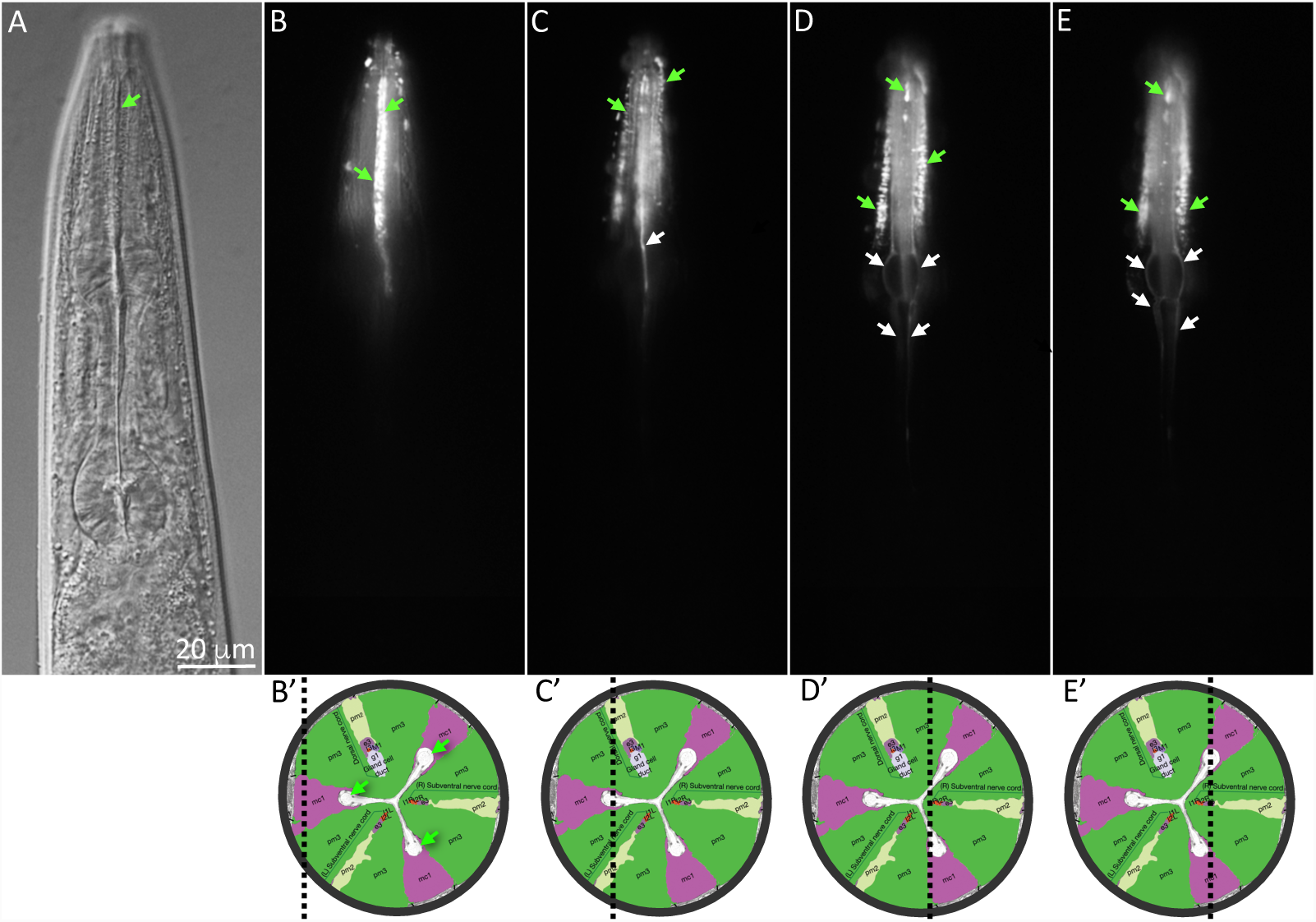
The Characterization of a Marginal Cell-Specific Promoter. **A.** A DIC image at the focal plane of the image shown in B. **B-E.** Four focal planes of an animal harboring an extra-chromosomal array with pPRHM1053, a construct consisting of a genomic copy of *pgp-14* (fosmid WRM64dE02) with YFP coding sequence inserted in frame at the C-terminus of PGP-14. PGP-14::YFP is expressed exclusively in the marginal cells of the anterior pharynx (corpus). White arrows indicate the basolateral surface of the marginal cells and green arrows point to the apical (luminal) surface of the cells that line the channels. **B’-E’.** Cartoons of cross sections of the pharynx corpus illustrating the approximate focal plane (black dotted line) of the corresponding images in B-E. Images are modified from Wormatlas. Marginal cells are purple; muscle is green, and the green arrows indicate the channels of the lumen.

**Figure S6.**
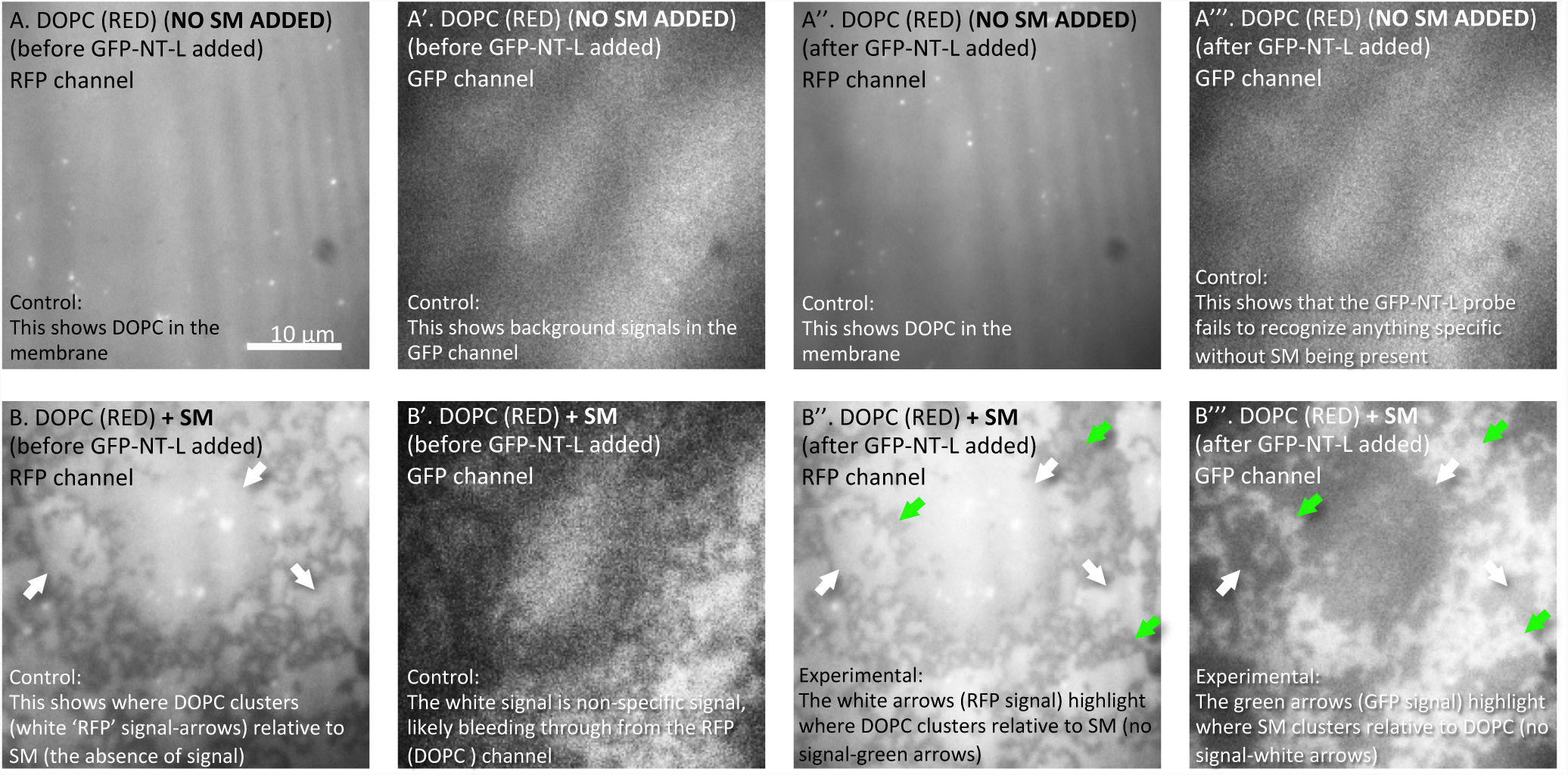
A Control Experiment that Shows that the GFP-NT-Lysenin Probe Binds Clustered Sphingomyelin. A-A’’’. The micrographs on the top row show a single lipid sample that is a suspended artificial 1,2-Dioleoyl-sn-glycero-3-phosphocholine (DOPC) lipid bilayer, either before (A and A’) or after (A’’ and A’’’) the GFP-NT-Lysenin probe was added to the sample. All samples shown are stained with Dil C, which binds DOPC and fluoresces red. Fluorescent signal is seen in the RFP channel (A and A’’), while only background fluorescence is seen in the GFP channels (A’ and A’’’). **B-B’’’.** The micrographs on the bottom row show a single lipid sample of DOPC mixed with sphingomyelin (SM) and otherwise treated in the same way as described above in A-A’’’. In the bottom row, homotypic clustering of the DOPC and SM lipids can be seen. DOPC clusters can be seen in the RFP channel (white arrows) (B and B’’) that are complementary to the GFP-NT-Lysenin fluorescent signal seen in the GFP channel (green arrows) after the GFP-NT-Lysenin probe (20µg/mL) has been added (15 minutes at room temperature) (B’’’). The lipid bilayers were prepared in 10 mM HEPES buffer, 150 mM NaCl, pH7.4 as previously described [41]. The scale is the same for all panels.

**Figure S7.**
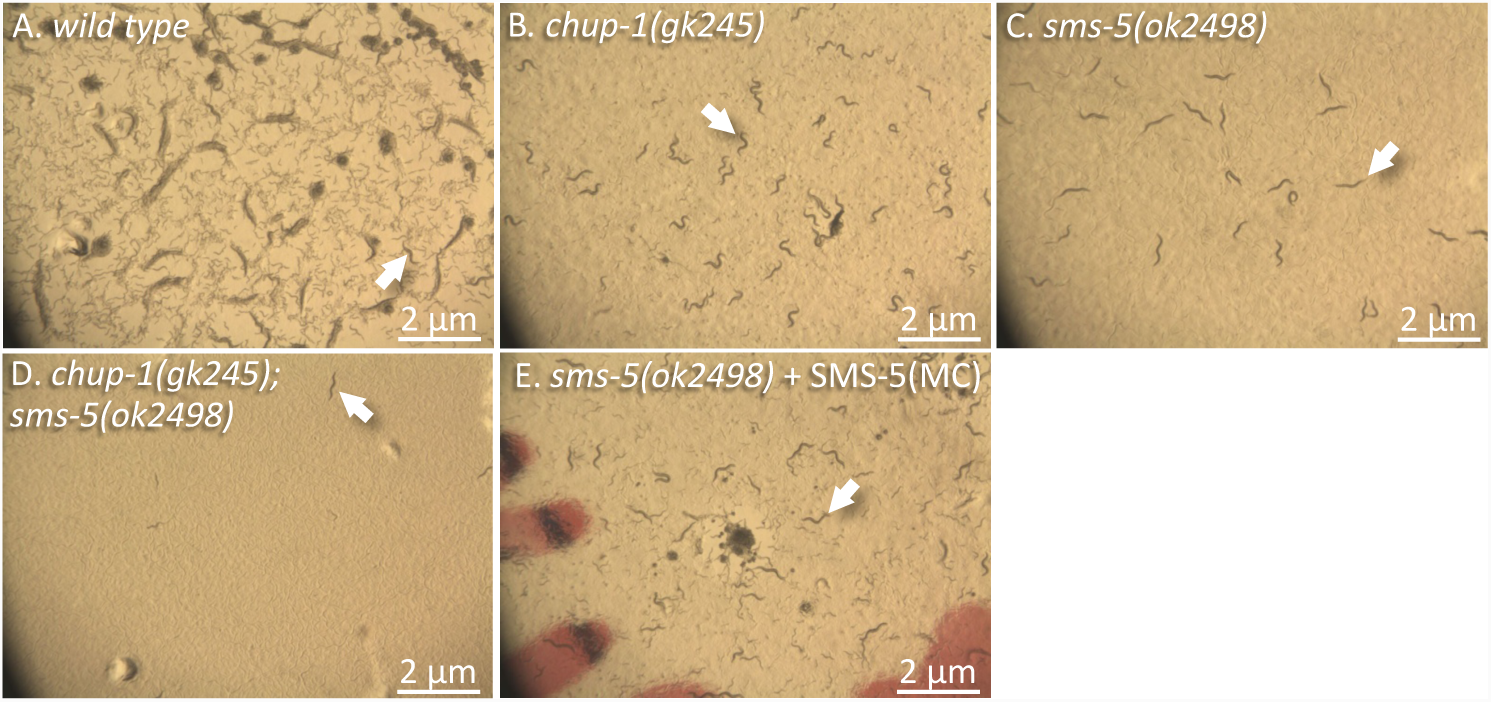
Examples of Cultures of Strains Grown in Cholesterol-Limited Conditions. The micrographs show a sector of the surface of the petri dish on which the indicated strains are grown in cholesterol(xol)-limited conditions (containing 50 ng/mL xol), corresponding to the data reported in Figure 8a. Arrows indicate representative adult animals on each plate. SMS-5(MC) represents marginal-cell specific expression of SMS-5 from the *trIs104* integrated transgenic array. The rust-colored streaks in the background of ‘E’ are part of the labeling of the plate. See methods for additional details.

**Figure S8.**
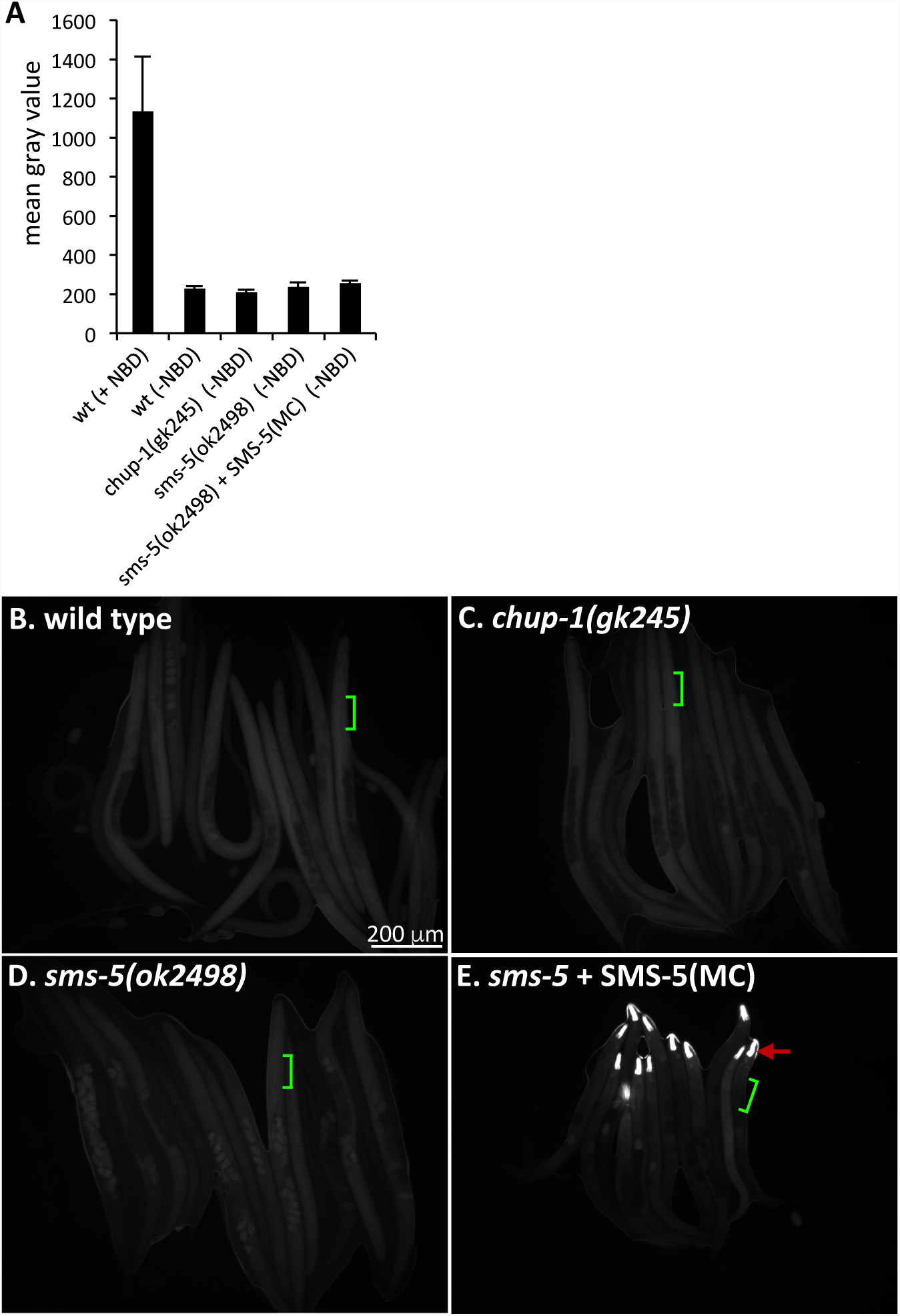
Measurement of Auto-Fluorescent Signal in Strains Incubated without NBD-cholesterol. **A.** Autofluorescent signal of the indicated strains incubated without NBD-cholesterol (-NBD) compared to wild type incubated with NBD-cholesterol (+NBD) using the same filter sets as that used to analyze the NBD-cholesterol signal in Figure 8. There are no significant differences in background signal intensity in any of the (-NBD) mutant strains compared to the wild type incubated without NBD-cholesterol (*p*>0.25). Standard error of the mean is shown. **B-E**. Images of multiple animals after the indicated strains were incubated without NBD-cholesterol for six days. The scale is the same for all images. Green lines exemplify the area used in each animal to calculate signal in ‘A’. Red arrow indicates the expression of the YFP marker used to mark the presence of the rescuing transgene *trIs104*. In ‘A’ and ‘E’, SMS-5(MC) represents marginal-cell (MC) specific expression of SMS-5 from the *trIs104* integrated transgenic array.

## Supplemental Datasets

**Supplemental Dataset 1.** An Excel file showing our survey of 238 wactive molecules for their ability to form objects in wild type (N2) L1 *C. elegans* larvae at a concentration of 30 μM. The file also contains the Chembridge Inc IDs, basic physicochemical properties, and a summary of the viability assays for all 238 compounds.

**Supplemental Dataset 2.** An excel file showing the results of our whole-genome sequence analyses of 17 wact-190-resistant mutants. The mutations associated with the 29 *pgp-14* alleles that we isolated will be published elsewhere.

## Cite Literature

1. Samuel, B.S., et al., Caenorhabditis elegans responses to bacteria from its natural habitats. Proc Natl Acad Sci U S A, 2016. 113(27): p. E3941–9.

2. Brenner, S., The Genetics of Caenorhabditis elegans. Genetics, 1974. 77: p. 71–94.

3. Riddle, D.L. A Genetic Pathway for Dauer Larvae Formation in Caenorhabditis elegans. in Stadler Geneti. Symp. 1977.

4. Kurzchalia, T.V. and S. Ward, Why do worms need cholesterol? Nat Cell Biol, 2003. 5(8): p. 684–8.

5. Hieb, W.F. and M. Rothstein, Sterol requirement for reproduction of a free-living nematode. Science, 1968. 160(3829): p. 778–80.

6. Frezal, L. and M.A. Felix, C. elegans outside the Petri dish. Elife, 2015. 4.

7. Avery, L. and B.B. Shtonda, Food transport in the C. elegans pharynx. J Exp Biol, 2003. 206(Pt 14): p. 2441–57.

8. Fang-Yen, C., L. Avery, and A.D. Samuel, Two size-selective mechanisms specifically trap bacteria-sized food particles in Caenorhabditis elegans. Proc Natl Acad Sci U S A, 2009. 106(47): p. 20093–6.

9. Albertson, D.G. and J.N. Thomson, The pharynx of Caenorhabditis elegans. Philos Trans R Soc Lond B Biol Sci, 1976. 275(938): p. 299–325.

10. Merris, M., et al., Sterol effects and sites of sterol accumulation in Caenorhabditis elegans: developmental requirement for 4alpha-methyl sterols. J Lipid Res, 2003. 44(1): p. 172–81.

11. Baiga, T.J., et al., Metabolite induction of Caenorhabditis elegans dauer larvae arises via transport in the pharynx. ACS Chem Biol, 2008. 3(5): p. 294–304.

12. Kwok, T.C., et al., A small-molecule screen in C. elegans yields a new calcium channel antagonist. Nature, 2006. 441(7089): p. 91–5.

13. Luciani, G.M., et al., Dafadine inhibits DAF-9 to promote dauer formation and longevity of Caenorhabditis elegans. Nat Chem Biol, 2011. 7(12): p. 891–3.

14. Burns, A.R., et al., Caenorhabditis elegans is a useful model for anthelmintic discovery. Nat Commun, 2015. 6: p. 7485.

15. Shaham, S., WormBook: Methods in Cell Biology, in WormBook, V. Ambros, Editor. 2006, The C. elegans Research Community.

16. Ostermeier, C., Antibody Fragment-Mediated Crystallization of Integral Membrane Proteins: A Review, in Methods and Results in Crystallization of Membrane Proteins, S. Iwata, Editor. 2003, International University Line: La Jolla, California. p. 73–86.

17. Matsuda, R., A. Nishikawa, and H. Tanaka, Visualization of dystrophic muscle fibers in mdx mouse by vital staining with Evans blue: evidence of apoptosis in dystrophin-deficient muscle. J Biochem, 1995. 118(5): p. 959–64.

18. Eckhardt, E.R., et al., Asymmetric distribution of phosphatidylcholine and sphingomyelin between micellar and vesicular phases. Potential implications for canalicular bile formation. J Lipid Res, 1999. 40(11): p. 2022–33.

19. Sharom, F.J., Flipping and flopping--lipids on the move. IUBMB Life, 2011. 63(9): p. 736–46.

20. Huitema, K., et al., Identification of a family of animal sphingomyelin synthases. EMBO J, 2004. 23(1): p. 33–44.

21. Tafesse, F.G., P. Ternes, and J.C. Holthuis, The multigenic sphingomyelin synthase family. J Biol Chem, 2006. 281(40): p. 29421–5.

22. Goni, F.M. and A. Alonso, Biophysics of sphingolipids I. Membrane properties of sphingosine, ceramides and other simple sphingolipids. Biochim Biophys Acta, 2006. 1758(12): p. 1902–21.

23. Mattei, B., et al., Membrane permeabilization induced by Triton X-100: The role of membrane phase state and edge tension. Chem Phys Lipids, 2017. 202: p. 28– 37.

24. Ohvo-Rekila, H., et al., Cholesterol interactions with phospholipids in membranes. Prog Lipid Res, 2002. 41(1): p. 66–97.

25. Ishitsuka, R., et al., A lipid-specific toxin reveals heterogeneity of sphingomyelin-containing membranes. Biophys J, 2004. 86(1 Pt 1): p. 296–307.

26. Yilmaz, N., et al., Molecular mechanisms of action of sphingomyelin-specific pore-forming toxin, lysenin. Semin Cell Dev Biol, 2018. 73: p. 188–198.

27. Hao, L., et al., Clozapine Modulates Glucosylceramide, Clears Aggregated Proteins, and Enhances ATG8/LC3 in Caenorhabditis elegans. Neuropsychopharmacology, 2017. 42(4): p. 951–962.

28. Valdes, V.J., et al., CUP-1 is a novel protein involved in dietary cholesterol uptake in Caenorhabditis elegans. PLoS One, 2012. 7(3): p. e33962.

29. Merris, M., et al., Long-term effects of sterol depletion in C. elegans: sterol content of synchronized wild-type and mutant populations. J Lipid Res, 2004. 45(11): p. 2044–51.

30. Matyash, V., et al., Distribution and transport of cholesterol in Caenorhabditis elegans. Mol Biol Cell, 2001. 12(6): p. 1725–36.

31. Holthuis, J.C., et al., The organizing potential of sphingolipids in intracellular membrane transport. Physiol Rev, 2001. 81(4): p. 1689–723.

32. Chachaty, C., et al., Building up of the liquid-ordered phase formed by sphingomyelin and cholesterol. Biophys J, 2005. 88(6): p. 4032–44.

33. Burns, A.R., et al., The novel nematicide wact-86 interacts with aldicarb to kill nematodes. PLoS Negl Trop Dis, 2017. 11(4): p. e0005502.

34. Burns, A., et al., High-Throughput Screening of Small Molecules for Bioactivity and Target Identification in Caenorhabditis elegans. Nat Protocols, 2006. 1: p. 1906–1914.

35. Tursun, B., et al., A toolkit and robust pipeline for the generation of fosmid-based reporter genes in C. elegans.PLoS One, 2009. 4(3): p. e4625.

36. Yamaji, A., et al., Lysenin, a novel sphingomyelin-specific binding protein. J Biol Chem, 1998. 273(9): p. 5300–6.

37. Maekawa, M., et al., Staurosporines decrease ORMDL proteins and enhance sphingomyelin synthesis resulting in depletion of plasmalemmal phosphatidylserine. Sci Rep, 2016. 6: p. 35762.

38. Ruvkun, G. and M. Finney, Antibody staining of formaldehyde-fixed worms., in WormAtlas, D. Hall and Z.F. Altun, Editors. 2012.

39. Hall, D.H., E. Hartwieg, and K.C. Nguyen, Modern electron microscopy methods for C. elegan. Methods Cell Biol, 2012. 107: p. 93–149.

40. Altun, Z.F. and D.H. Hall, Alimentary System, Pharynx. WormAtlas, 2009.

41. Oreopoulos, J. and C.M. Yip, Probing membrane order and topography in supported lipid bilayers by combined polarized total internal reflection fluorescence-atomic force microscopy. Biophys J, 2009. 96(5): p. 1970–84.

